# Biological age prediction using a novel DNN model based on steroid metabolic pathways

**DOI:** 10.1101/2024.11.07.622443

**Authors:** Qiuyi Wang, Zi Wang, Kenji Mizuguchi, Toshifumi Takao

**Author notes:** Corresponding author. (Z. W.); (T. T.). These authors contributed equally to this work (Qiuyi Wang, Zi Wang). Shimadzu Corporation, Kyoto 604-8511, Japan (Q. W.).

## Abstract

Aging involves the progressive accumulation of cellular damage, leading to systemic decline and age-related diseases. Despite advances in medicine, accurately predicting Biological Age (BA) remains challenging due to the complexity of aging processes and the limitations of current models. This study introduces a novel method for predicting BA using a Deep Neural Network (DNN) based on steroid metabolic pathways. We analyzed 22 steroids from 148 serum samples of individuals aged 20 to 73, using 98 samples for model training and 50 for validation. Our model reflects the often-overlooked fact that aging heterogeneity expands over time and uncovers sex-specific variations in steroid interactions. This study identified key markers, including cortisol (COL), which underscore the role of stress-related and sex-specific steroids in aging. The resulting model establishes a biologically meaningful and robust framework for predicting BA across diverse datasets, supporting more targeted strategies in aging research and disease management.

## Introduction

Aging is a complex and inevitable process involving the accumulation of cellular and molecular damage, leading to functional decline and an increased risk of age-related diseases (*1, 2*). Conditions such as Alzheimer’s disease, Parkinson’s disease, and osteoporosis are closely tied to aging and contribute significantly to the health challenges faced by the elderly (*2-6*). Despite medical advancements, these diseases remain incurable, with current strategies focused on slowing their progression through early diagnosis and management (*7-10*). Accurately assessing an individual’s Biological Age (BA), which reflects their physiological state, is essential for understanding aging and developing effective interventions. Unlike Chronological Age (CA), which simply measures the passage of time, BA provides insights into the biological processes underlying aging (*11*). However, determining BA is complex, as it is influenced by both genetic and non-genetic factors, and no universally accepted standards for BA measurement currently exist (*12-17*). Early methods, which used phenotypic indicators like lung capacity and grip strength, lacked precision and standardization, limiting their predictive utility for aging-related diseases (*18-23*).

In recent years, researchers have shifted from surface-level indicators to more intrinsic measures that better capture physiological aging. Common diagnostic tools like complete blood counts and biochemical tests are frequently used to model BA, offering valuable insights into an individual’s health (*24*). However, these markers often fail to provide a direct window into the specific physiological or metabolic pathways that drive aging. To address this, omics technologies—such as genomics, epigenomics, transcriptomics, proteomics, and metabolomics—have been employed to analyze aging at a molecular level (*25*). These approaches generate high-dimensional data, revealing complex interactions among potential biomarkers. Given the significant role of non-genetic factors in aging, methods like epigenomics and metabolomics, which are sensitive to environmental and lifestyle influences, have proven particularly effective in enhancing the accuracy of BA models (*12, 13, 26*). Building on these advancements, steroid hormones have emerged as crucial indicators of physiological aging due to their regulation of key metabolic processes (*27-30*). Stress-related corticosteroids and sex hormones, both of which strongly correlate with aging, present a promising avenue for refining BA predictions. By incorporating interactions between these hormones, BA models can more accurately reflect the underlying physiological state of aging individuals.

Developing precise BA models has become a central focus in bioinformatics, with researchers using various biomarkers to estimate BA (*31*). Methods such as Least Absolute Shrinkage and Selection Operator (LASSO) and Ridge regression have been applied to DNA methylation and proteomics data (*32-34*). While these methods are effective at identifying linear relationships, they often overlook the biomarkers linked to metabolic pathways, which are critical to understanding aging (*35, 36*). Traditional machine learning techniques, though useful for preventing overfitting and balancing model complexity, struggle to capture the non-linear interactions inherent in biological systems (*12*). As a result, these methods may miss the intricate biological processes underlying aging and fail to account for the substantial impact of environmental and lifestyle factors.

To overcome these challenges, modern machine learning techniques—such as Support Vector Machines (SVMs) (*37, 38*), Random Forests (*39*), and Deep Neural Networks (DNNs) (*40-42*) —have gained prominence. These approaches excel at modeling non-linear relationships, making them particularly well-suited for capturing the complex biological processes involved in aging. DNNs, in particular, are effective at handling high-dimensional data and are widely used for tasks such as predicting BA. Researchers, including Levine, Mamoshina, and Putin, have employed public datasets of blood tests and biochemical markers to predict BA using DNNs, leveraging their capacity for feature learning (*40-42*). Mamoshina also applied similar techniques to gene expression data from muscle samples, identifying age-related markers (*43*). Despite their fitting capabilities, however, DNNs are prone to overfitting, especially when involving numerous hidden layers and nodes, which can reduce performance on unseen data (*44-46*). Regularization techniques, cross-validation, and data augmentation are typically employed to mitigate these issues, but challenges remain. One major limitation of current BA models is their emphasis on minimizing prediction errors—often measured by Mean Absolute Error (MAE) or Mean Squared Error (MSE)—which may not fully capture the increasing heterogeneity of aging over time (*47, 48*). Additionally, DNNs often function as “black boxes,” making it difficult to derive biological meaning from the learned features. Addressing the issues of aging heterogeneity, biological interpretability, and feature selection remains a pivotal area of research.

In this study, we developed a DNN model centered on steroid metabolic pathways to enhance the accuracy of BA prediction. Steroids were quantified using an *in-house* liquid chromatography–tandem mass spectrometry (LC-MS/MS) method (*49*), with the resulting data stratified into four groups according to sex and designation for training or independent validation (Fig. 1A). To address physiological and experimental variability, we applied tailored data scaling techniques that preserved the inherent steroid ratios while achieving reliable alignment between the training and validation datasets (Fig. 1B). The DNN model also incorporates a custom-designed loss function, specifically constructed to account for the progressive heterogeneity of aging—a feature largely neglected in previous predictive models (Fig. 1C). Furthermore, the DNN architecture was structured to capture interactions within key steroid pathways, thus substantially enhancing the model’s biological interpretability (Fig. 1D). Through consideration of sex-specific steroid interactions and validation with independent datasets, our goal was to establish a DNN-based BA prediction model that effectively represents diverse aging patterns across populations and reflects fundamental biological pathways.

**Fig. 1.**
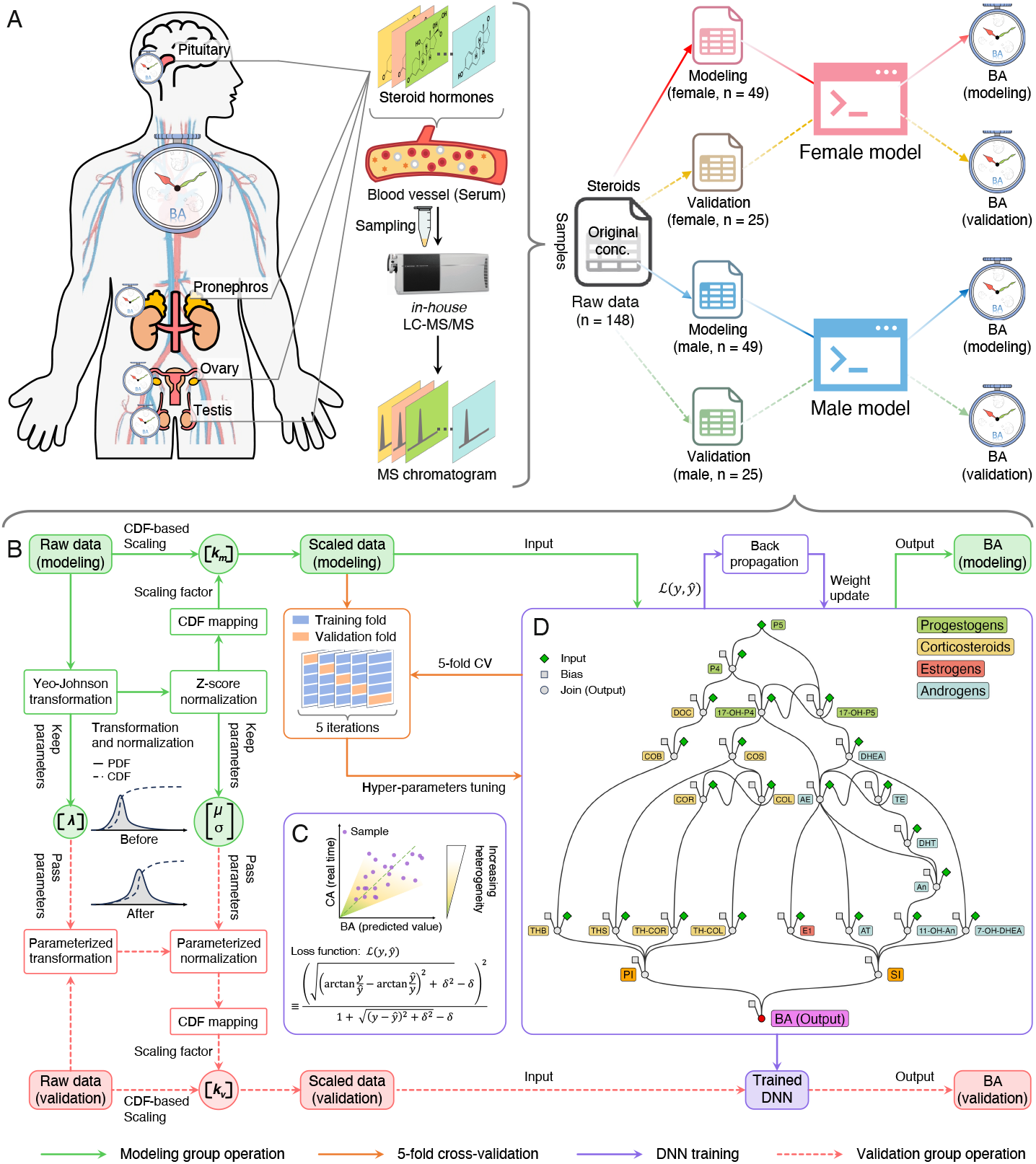
Pathway-based DNN model for BA prediction from serum steroid profiling via LC-MS/MS. (**A**) Steroid hormone quantification in blood using the LC-MS/MS method, with data divided into four groups by sex and assigned to either training with 98 samples or validation with 50 samples. (**B**) Tailored data scaling techniques were applied to address physiological and experimental variability, ensuring a reliable training dataset. This process included a CDF-based proportional scaling method, Yeo-Johnson transformation, and z-score normalization to maintain relative steroid concentration differences across samples. (**C**) The DNN model reflects aging heterogeneity and lifestyle-related variations by integrating known steroid pathways. A custom WSATL function balances prediction accuracy by weighting differences between BAs and CAs, preventing overfitting. (**D**) The DNN model identified key steroid pathways associated with aging, enhancing biological interpretability for BA prediction. The framework, based on steroid metabolic pathways extracted from the KEGG database, consists of 25 nodes connected by 37 directed edges, revealing the interactions that influence aging.

## Results

### DNN dataset generation via steroid quantification using LC-MS/MS

We developed a method to quantify 30 steroid hormones in serum, with the list of compounds and their structures shown in Table S1 and Fig. S1 and the experimental parameters outlined in Table S2. Validation results, including assessments of limit of quantitation (LOQ), linearity, recovery, precision, and accuracy (Table S3), confirm the method’s robustness for steroid quantification.

We used this validated method to quantify 22 steroids in 150 individuals, aged 20 to 73, with detailed results presented in Table S4. Out of the 100 samples used for modeling, two were excluded due to one exceeding the maximum LOQ and the other having most compounds below the LOQ, while 50 samples were utilized for validation (Fig. 1A). As shown in Fig. S2 and Table S5, the concentrations varied widely but mostly aligned with previous studies. Differences in estrone (E1) levels in female samples likely stemmed from menstrual cycle variations. The broader range of 7α-hydroxydehydroepiandrosterone (7-OH-DHEA) in our study may reflect the inclusion of younger, more diverse participants, whereas prior research focused on older individuals (aged 50 to 91). Comparisons with previous studies on tetrahydrocortisol (TH-COL), tetrahydrocortisone (TH-COR), 11-β-hydroxyandrosterone (11-OH-An), tetrahydrocorticosterone (THB), tetrahydrodeoxycortisol (THS), and adrenosterone (AT) are limited due to smaller sample sizes in those studies (fewer than 20 subjects), while our dataset is more robust (see Table S6).

To conduct DNN modeling, we gathered additional information about the individual subjects, including demographic labels such as CA, sex, ethnicity, ABO blood types, Rh blood types, and smoking habits (applicable only to the independent validation datasets). Due to individual variability among different subjects in this dataset, appropriate scaling of the steroid concentrations is required.

### Maintaining biological consistency and minimizing batch variability

Building on the initial observations of raw steroid concentration data, we implemented a cumulative distribution function (CDF)-based proportional scaling method to refine the dataset, aiming to preserve inherent steroid concentration ratios while reducing batch variability across samples. This approach transforms each sample’s steroid concentrations by aligning them with a reference distribution, facilitating consistent downstream modeling. Specifically, the scaling process began with a Yeo-Johnson transformation followed by z-score normalization to approximate normality, standardizing across variables while retaining relative concentration differences.

The similarity in distribution patterns for scaled concentrations in both the modeling and validation datasets indicates that scaling effectively preserves the relative steroid concentration ratios within each sex (Fig. 2A). Notably, proportional differences among steroids are maintained, with reduced inter-group variability, as shown by the closer alignment of concentration distributions across cohorts compared to the original data.

**Fig. 2.**
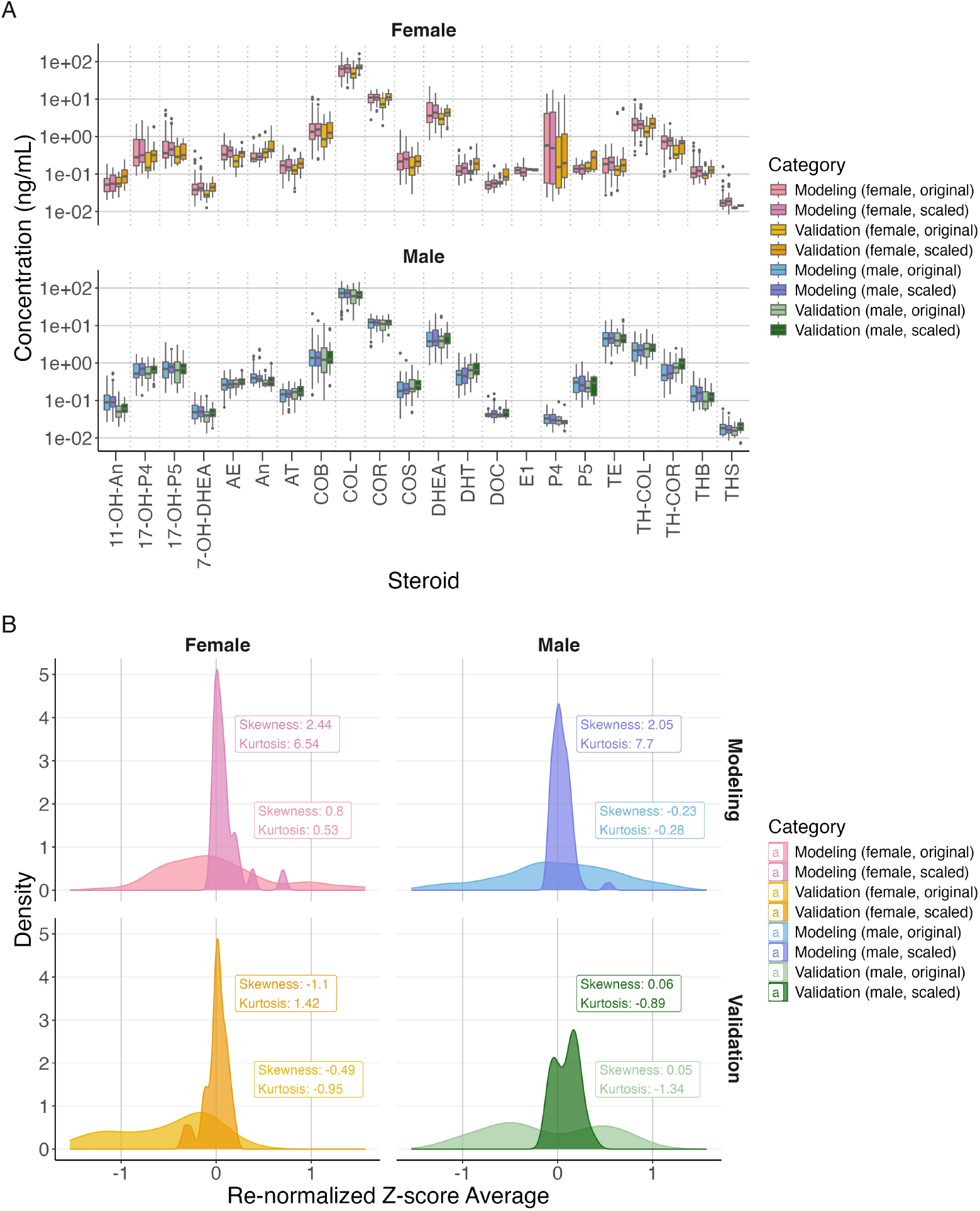
Impact of CDF-based proportional scaling on steroid concentration distributions. (**A**) Steroid concentration distributions before and after scaling for modeling and validation datasets, stratified by sex. (**B**) Density distributions of Z-scores for original and scaled steroid concentrations across modeling and validation datasets by sex.

To further evaluate scaling’s impact on overall sample distribution, we re-normalized the scaled steroid concentrations into z-scores for each individual and averaged these values across all steroids. This was then compared with z-scores derived from the original, unscaled steroid concentrations to assess cumulative profiles (Fig. 2B). In both the modeling and validation groups, the distribution shifted from a broad, flat pattern in the original data to a more concentrated distribution centered around a z-score of zero, indicating reduced sample-to-sample variability. This outcome suggests that scaling effectively aligns individual distributions and minimizes batch effects while preserving biologically relevant concentration gradients between steroids.

Moreover, analysis of the scaled steroid concentration distributions in the modeling dataset showed no significant differences across ABO blood types (Fig. S3A), Rh blood types (Fig. S3B), or ethnicities (Fig. S4), further supporting the robustness of the scaling method. This finding suggests that these demographic labels are unlikely to influence subsequent modeling outcomes. Together, these results underscore the dual advantages of this CDF-based proportional scaling approach: preserving essential steroid concentration patterns and minimizing biases that could otherwise compromise model accuracy and generalizability. The enhanced uniformity across samples and consistency in steroid ratios within the scaled data are expected to strengthen model robustness, ensuring that input data accurately reflects biologically relevant variation.

### DNN design: unveiling pathway biological features and sex-specific insights

Building on the achieved uniformity and minimized batch variability in scaled steroid concentrations across demographic subgroups, we implemented our metabolic pathway-based DNN to predict BA. The model’s architecture is explicitly designed to reflect the sequential stages of steroid biosynthesis: starting from pregnenolone (P5) as the precursor, progressing through intermediate metabolites, and culminating with the physiological indices, Pressure Index (PI) and Sexual Index (SI), along with the final BA prediction. This structured design enables the DNN to model steroid pathway interactions known to influence aging, particularly under various stress and hormonal conditions.

To embed biologically meaningful interactions, we initialized the edge weights according to Spearman correlations among steroids and between steroids and CA (Fig. S5). This initialization avoids random weights—a common source of instability in DNN models—and reduces biases linked to irrelevant biological processes by uniformly setting initial bias values to zero. To capture the heterogeneity of aging across CA, we employed a custom Weighted Symmetric Arc-Tangent Loss (WSATL) function, which penalizes disproportionate predictions and maintains symmetry in the model’s handling of high and low biases across different CA ranges. This approach accounts for the progressive variability in aging, a complexity often overlooked in traditional models.

Training optimization was conducted through 5-fold cross-validation to select hyperparameters, specifically the learning rate (*lr*) and number of epochs (*t*), ensuring a balance between validation fold loss, training stability, and computational efficiency (Fig. S6). Based on these evaluations, a learning rate of 0.005 with 4000 epochs was selected for females, and a learning rate of 0.003 with 8000 epochs for males, yielding smooth convergence with minimal loss fluctuation across iterations and contributing to stable training dynamics (Fig. S7A, B). The scatter distribution of predicted BA against actual CA reflects the intended design of the loss function, illustrating that the heterogeneity of BA expands over time and that both overestimations and underestimations across various CA segments are balanced (Fig. S7C).

The final trained models, depicted in Fig. 3A (female) and Fig. 3B (male), illustrate the interaction patterns across steroid pathways, highlighting the nodes and connections with the greatest impact on BA predictions (Table S7). Visualization of the scaled weights between pathway components (connection weights) enables the identification of a hierarchy of steroid influence on BA prediction. The average contribution of each node (node influence) propagates through the pathway network, providing an indication of each node’s relative role in predicting BA. Additionally, categorizing nodes by origin (component type) allows for the differentiation of endogenous and exogenous influences, offering insights into their respective contributions within the model.

**Fig. 3.**
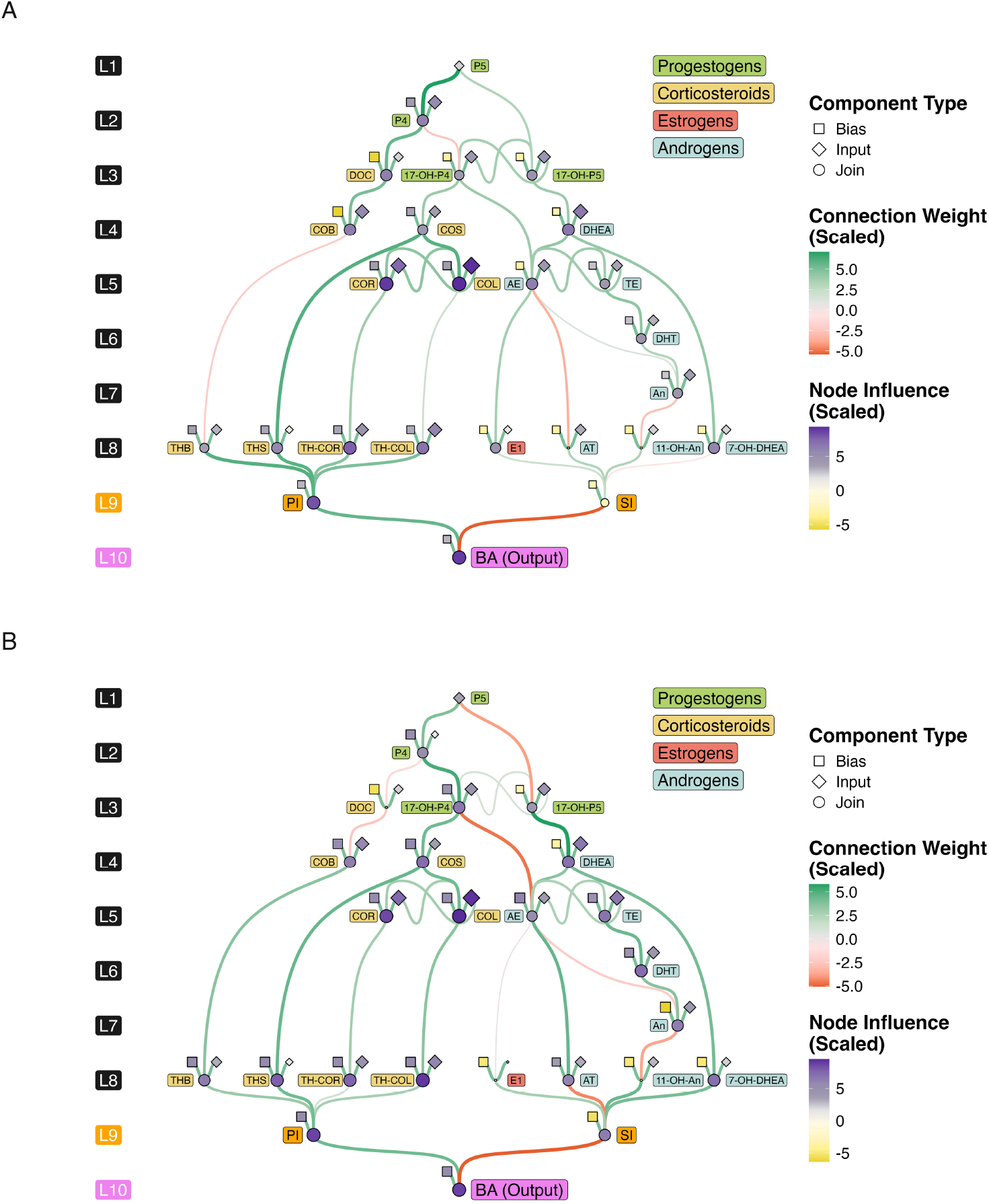
Visualization of the DNN model constructed on steroid metabolic pathways. Sex-specific variations in steroid pathways for female (**A**) and male (**B**) models. Distinct colors are used to represent different steroid classes in the steroid labels. Connection weights indicate the hierarchical interactions of steroid influences on BA prediction. Node influence reflects the average contribution of each node as it propagates through the pathway network. Component types illustrate the various sources of endogenous and exogenous influences. Detailed edge weight values and node values can be found in Tables S7. Bias, contribution from external pathways; Input, initial concentration; Join, summarized contributions from upstream metabolites.

Notably, corticosteroid and sex hormone pathways significantly contribute to BA, with distinct impacts observed between female and male models, consistent with physiological differences. Corticosteroid nodes show strong positive associations with PI in both models, aligning with established research linking elevated corticosteroids to stress-related aging effects and supporting the hypothesis that stress pathways play a substantial role in aging across sexes. The DNN also captured sex-specific patterns within the steroid pathway: estrogen-related nodes, such as the E1 join node, exhibited heightened influence in the female model, while androgen-related nodes, such as the AT join node, were more pronounced in the male model. This finding emphasizes the physiological specificity embedded within the DNN, in line with sex-specific hormonal profiles and their aging implications.

These pathway-based DNN models offer a robust, biologically informed approach to predicting BA by precisely leveraging steroid interactions. To validate the interaction patterns identified, we assessed the model’s predictive accuracy and generalizability, particularly in capturing BA variability across diverse independent validation datasets.

### DNN model performance and smoking impact on BA prediction

To assess the performance of the established DNN model, we analyzed the scatter distribution of predicted BA against actual CA for both the model training group and the independence validation group. The distribution of individual prediction results suggests that the boundaries of a 2-fold change can be interpreted as physiological thresholds indicative of a younger or older biological state. Notably, the majority of predictions fall within this 2-fold change range (Fig. 4A). Statistical analysis of the WSATL value across the various groups reveals no significant differences in prediction losses, indicating consistent performance across the cohorts (Fig. S8).

**Fig. 4.**
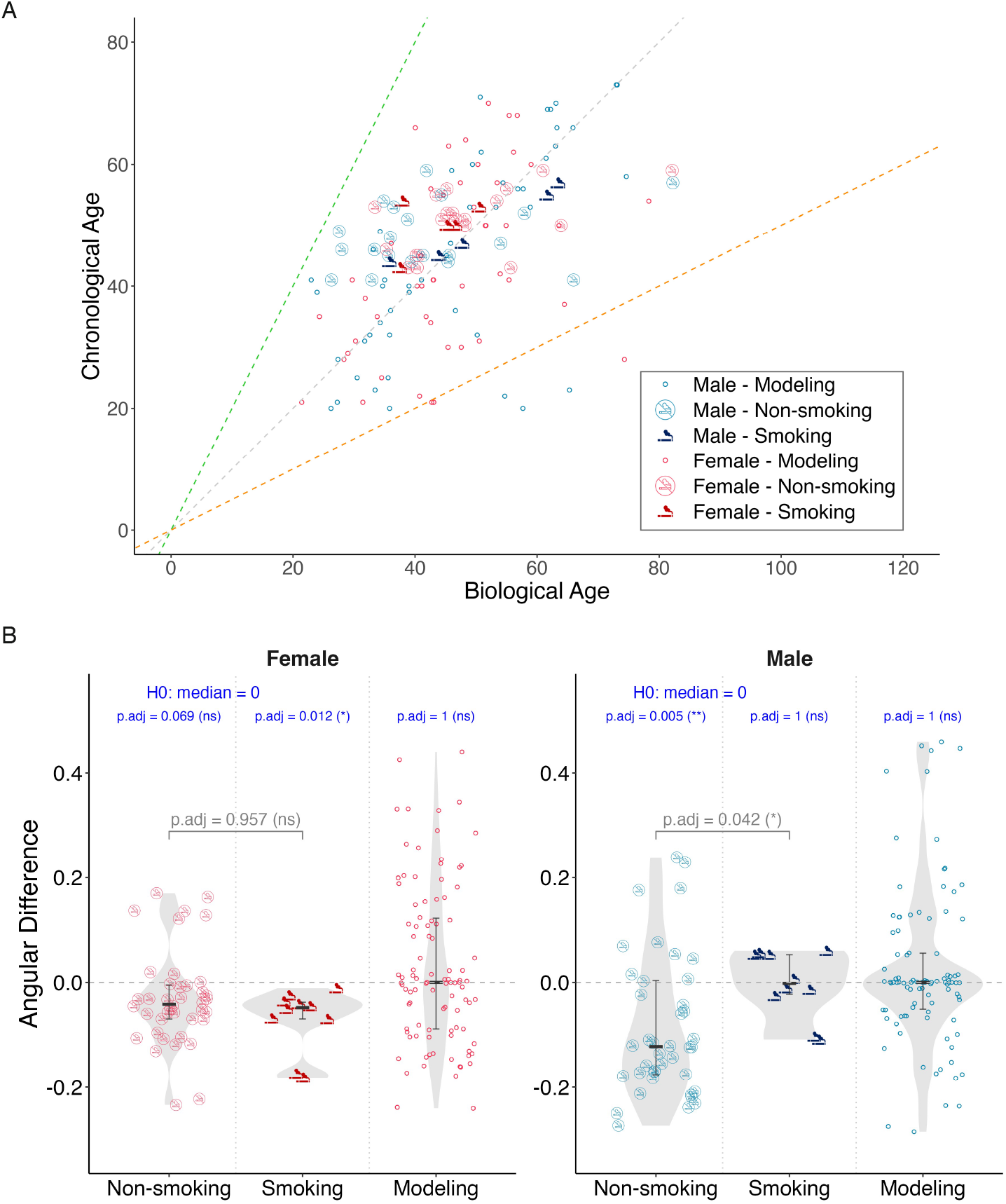
Performance of the DNN model and smoking impact on BA prediction. (**A**) Scatter plot of predicted BA versus CA for modeling and validation samples, including both smoking and non-smoking groups. The dashed lines represent the boundaries of the two-fold change, which can be interpreted as physiological thresholds indicative of a younger or older biological state. (**B, C**) Statistical analysis of angular differences for female (**B**) and male (**C**) samples. The gray p-values indicate the differences between the smoking and non-smoking groups, while the navy values represent the p-values for each group in relation to the null hypothesis (H0) set to a median value of zero. Statistical comparisons were performed using the Wilcoxon test adjusted by Bonferroni correction. The groups include non-smoking (validation, n = 40), smoking (validation, n = 10) for each sex, and modeling (smoking status unknown. female, n = 48; male, n = 49) individuals. ns P_adj_ ≥ 0.05; * P_adj_ < 0.05; ** P_adj_ < 0.01.

Additionally, the validation group dataset includes information regarding smoking habits, allowing us to further investigate the impact of smoking on BA predictions. By examining the angular difference (*φ*) among individuals, we find that the smoking subgroup of females shows no significant difference in *φ* compared to their non-smoking counterparts (Fig. 4B). In contrast, the smoking subgroup of males exhibits a statistically significant difference in *φ* when compared to non-smokers (Fig. 4C), suggesting that smoking habits may accelerate biological aging in male individuals (*50, 51*).

It is important to note that while the modeling group lacks explicit smoking habit labels, which obscures the absolute positioning of the reference frame, the relative distribution of *φ* among different smoking habits in the validation group remains discernible. Specifically, when using the model group data as a baseline, the test of *φ* against a value of zero shows no significant differences across both male and female model groups. However, the relative distribution differences in *φ* among the validation group’s smoking habits are preserved, highlighting the robustness of our DNN model in capturing the nuanced effects of lifestyle factors on biological aging.

### Sensitivity analysis for identifying aging key markers

To assess the sensitivity of the established DNN model, we examined the impact of doubling the input values of each steroid node on the output values of the BA node (Fig. 5A). Notably, both the female and male DNN models revealed that cortisol (COL), a steroid associated with stress and present in relatively high concentrations, exerted a significant positive sensitivity effect on BA predictions, exceeding 40%. Additionally, in the female model, steroids such as 17α-hydroxyprogesterone (17-OH-P4), cortisone (COR), 11-deoxycortisol (COS), and TH-COL also demonstrated a stable positive influence on BA. In the male model, P5 and testosterone (TE) exhibited similar trends.

**Fig. 5.**
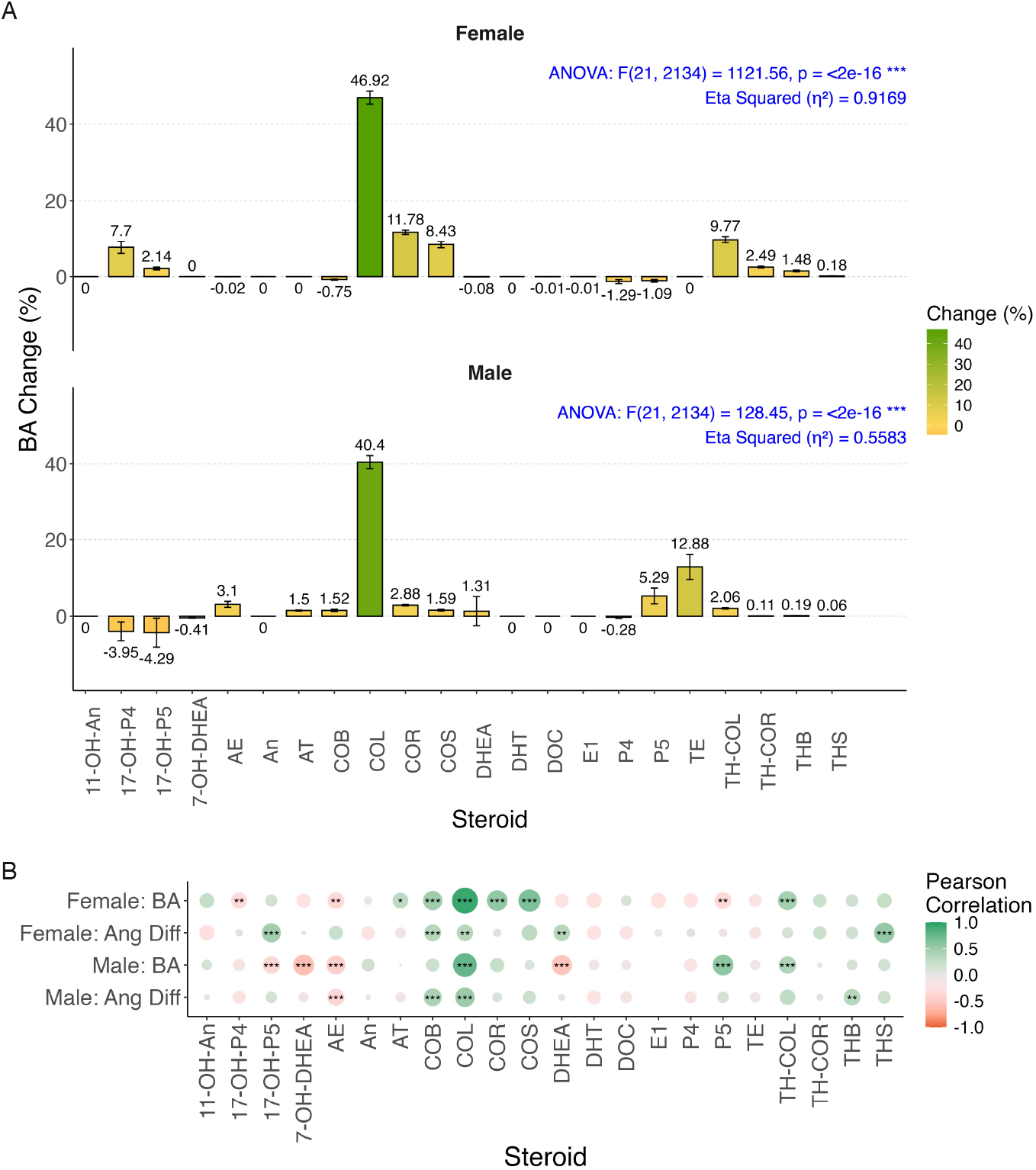
Sensitivity analysis and aging marker identification. (**A**) Percentage change in BA in response to a two-fold increase in each of the 22 steroid input nodes for sex-specific DNN models. Error bars represent the variability in sensitivity results across individuals. (**B**) Heatmaps of Pearson correlation coefficients between the 22 steroid input nodes and BA, as well as the angular difference between BA and CA in both male and female models. Pearson correlation test was assessed with Bonferroni correction. ns P_adj_ ≥ 0.05; * P_adj_ < 0.05; ** P_adj_ < 0.01; *** P_adj_ < 0.001.

Analysis of Variance (ANOVA) results indicated that the input values of the 22 steroids explained a substantial portion of the BA prediction model, achieving an explanatory ability (*η*^2^ value) of 0.9169 for females and 0.5583 for males, highlighting the reliability of our biological process modeling. Conversely, we did not identify any steroids with a consistent negative impact on BA, suggesting that the physiological regulation required to delay biological aging is more likely associated with the reduction of stress-related hormones.

Furthermore, an analysis of the scaled values of each steroid concerning smoking habits revealed no significant differences across sexes (Fig. S9), underscoring the necessity of considering the biological interactions among steroids. When evaluating BA as an absolute indicator of aging, alongside the angular difference (*φ*) of BA against CA as a relative aging indicator, COL demonstrated a strong linear correlation and high confidence for both indicators across gender models (Fig. 5B). This suggests that COL may serve as a robust marker for reflecting BA, with its associated physiological pathways likely containing important factors or processes related to aging regulation.

## Discussion

In this study, we developed a novel DNN model based on steroid metabolic pathways to predict BA. The model effectively captured the increasing heterogeneity of aging over time and the complex biological processes influenced by steroid interactions. After training the model on a well-structured cohort, we analyzed the intricate relationships between specific hormones and physiological aging processes. The DNN architecture allowed for a detailed examination of how different steroids impact BA, revealing notable sex-specific differences between male and female models. These findings underscore the distinct metabolic interactions in each sex and their influence on aging trajectories.

One strength of our model lies in its use of a CDF-based proportional scaling that preserves intrinsic steroid concentration ratios while minimizing variations due to experimental batch effects and individual total steroid levels. Although not central to the model’s design, this scaling approach contributed to the robustness and accuracy of predictions, which were validated with independent datasets. The model’s adaptability to more diverse datasets further underscore its potential for continuous refinement as additional data become available.

A significant finding from our analysis was the identification of several steroid markers associated with aging, with COL emerging as a particularly prominent marker. While COL is abundant and easily measurable, its contribution to aging in our model was treated as a linear factor. However, it is important to recognize that COL might serve as an indicator of aging rather than a direct cause, and its involvement in processes like gluconeogenesis and inflammation suppression requires further investigation into its upstream and downstream pathways. Additionally, many steroids, including COL, exhibit circadian variations, raising questions about the need for correcting these fluctuations in future models (*52, 53*).

When considering lifestyle factors, smoking status emerged as a notable variable, though our validation cohort lacked detailed information on other behaviors such as alcohol consumption and diet (*54-59*). Our results revealed that only male smokers exhibited a more accelerated aging trajectory compared to non-smokers. We hypothesize that this disparity may be due to the generally higher smoking frequency among males. In contrast, the lower smoking frequency in females, combined with unmeasured lifestyle factors that influence BA, may have obscured the aging effects in female smokers. Future studies with larger cohorts and more comprehensive lifestyle and physiological data will be crucial to further elucidate these relationships.

Despite the valuable insights provided by this study, several limitations should be acknowledged. The relatively small sample size and the lack of detailed lifestyle data across certain cohorts limit the generalizability of our findings. Additionally, our treatment of steroids in a static manner, without fully accounting for their broader biochemical pathways and circadian variations, could introduce biases. To address these limitations, future research should focus on longitudinal studies that track individuals over time and incorporate more dynamic approaches that capture steroid fluctuations. Expanding the dataset to include a wider range of environmental and behavioral factors, alongside deeper exploration of sex-specific metabolic pathways, will further enhance the predictive power and clinical relevance of the model. These improvements could help make the BA prediction model a valuable tool in personalized aging interventions, facilitating the identification of biomarkers and enabling more targeted strategies to modulate aging processes.

## Materials and Methods

### Chemicals

This study utilized thirty steroid standards and fourteen internal standards, as summarized in Fig. S1 and Table. S1. HPLC-grade methanol (MeOH), acetonitrile (ACN), and 99.998% trace metals basis lithium chloride (LiCl), as well as analytical reagent-grade acetic acid (AcOH), were purchased from Sigma-Aldrich (USA). HPLC-grade formic acid was obtained from FUJIFILM Wako Pure Chemical Corporation (Japan). Ultrapure water was produced using an Organo Puric ω system (Japan).

### Sample acquisition and cohort

Serum samples from 150 healthy individuals were obtained from BIOIVT (New York, US). All samples were collected in the US, imported to Japan on dry ice, and stored at -80 °C until analysis. The storage and study protocols were approved by the Institute for Protein Research’s ethics committee. Our study modeled BA using data from 100 healthy participants (50 females and 50 males, aged 20–73 years). The model was then applied to a validation cohort of 50 participants (25 females and 25 males, aged 40–59 years). Of the validation set, 40 participants were non-smokers, while 10 were smokers, each smoking at least 10 cigarettes per day.

### Sample preparation

Serum (240 µL) and 4.8 µL of isotope-labeled internal standard solution were added to a 1.5 mL tube (Eppendorf, Germany) and mixed, followed by the addition of 480 µL of ACN. The mixture was vortexed at 3,200 rpm for 30 seconds and incubated at 4 °C for 30 minutes, followed by centrifugation at 20,000 × g at 4 °C for 15 minutes to precipitate proteins. The supernatant was transferred to a 15 mL tube and diluted with 4.08 mL H_2_O, achieving a final acetonitrile concentration of 10% (v/v). After a second centrifugation at 19,000 × g at 4 °C for 15 minutes, the supernatant was applied to a Bond Elut C18 column (Agilent, USA), preconditioned with 80% ACN and 10% ACN. The sample was washed with 10% ACN and eluted with 1 mL of 80% ACN, which was collected in a new tube. The eluate was dried using a speed-vac and the residue redissolved in 24 *µL* of 40% MeOH. After centrifugation at 20,000 × g for 15 minutes, the supernatant was transferred, and analyzed by LC-MS/MS.

### LC-MS/MS analysis

LC-MS/MS quantification of steroids was performed using an Agilent 1290 Infinity II and a 6470 triple quadrupole mass spectrometer (Agilent) in positive ion mode, utilizing the multiple reaction monitoring (MRM) method. Chromatographic separation was achieved with an C18 column (Eclipse Plus C18 RRHD 2.1×100 mm, 1.8 μm, Agilent, USA) at 40 °C. The details of this method are reported in our previous method.

### CDF-based proportional scaling

To achieve cross-batch alignment while maintaining within-group proportionality, we calculated a scaling factor *k* for each individual dataset. This process began by applying the Yeo-Johnson transformation (*60*) *X*′ = *f*_YJ_(*X, λ*), to each variable *X* (representing the original concentration of steroids) to address non-normality. Here, *X*′ represents the transformed data, *f*_YJ_ is the transformation function:

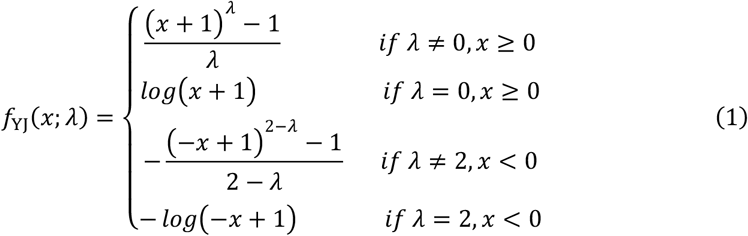

Here, *λ* was optimized for each variable (steroid) within the modeling group and retained as a parameter for transformations on independent validation data. Next, to standardize across individuals, we computed the z-score as:

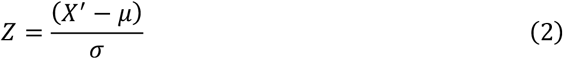

where *μ* is the mean and *σ* the standard deviation of *X′* within each steroid, both optimized within the modeling group and retained for parameterized normalization on independent validation data. Thus, for independent validation data, transformations and normalizations in Eq. (1) and Eq. (2) were performed using the retained parameters *λ, μ*, and *σ* from the modeling group. The z-scores *Z* were then mapped to a common cumulative distribution function (CDF) to derive the scaling factor *k*:

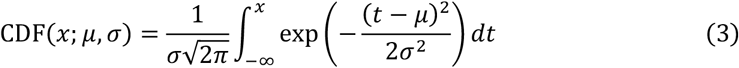

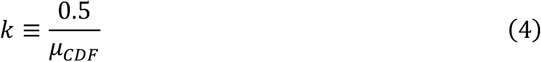

In Eq. (3), *μ* is the mean of the distribution, and *σ* its standard deviation. In Eq. (4), *μ*_*CDF*_ is the mean CDF value across all steroids for each individual, with 0.5 representing the cumulative distribution at Z = 0. Finally, each individual’s raw dataset was rescaled by *k* to achieve aligned concentration values:

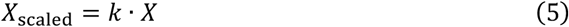

Here, *X*_scaled_ denotes the final scaled concentration, preserving proportional consistency across batches.

### Metabolic pathway-based DNN architecture

We extracted steroid-associated metabolic pathways from the KEGG database (*61, 62*) and encoded each pathway as input to a deep multilayer perceptron (DNN), designed to simulate steroid synthesis and metabolic processes in organisms. The DNN model was built with 25 components, beginning with P5, continuing through 21 intermediate steroids, and concluding with pressure index (PI) and sexual index (SI) outputs, with a final output for BA. To enhance biological interpretability, the model incorporated 22 input nodes representing scaled data for P5 and the 21 intermediate steroids, distributed across 8 layers. It also included 24 bias nodes (including PI, SI, and BA) to capture influences from other biological processes. These input and bias nodes collectively fed into 24 join nodes, which represent intermediate outputs for the 21 steroids, PI, and SI, as well as the final BA output. A total of 37 weighted edges linked upstream input or join nodes to downstream ones, capturing interactions among these biological components.

### Model loss function designing

WSATL function was designed to address the expanding heterogeneity of aging over time:

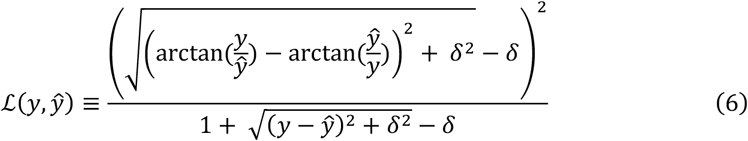

where *y* represents the true CA, and *ŷ* the predicted BA. The parameter *δ* (default set to 0.001) smooths the function 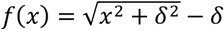, mitigating sharp deviations. The angular difference between 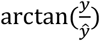 and 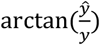 captures proportional similarity between BA and CA, while the denominator in Eq. 6 acts as a weighting factor, imposing greater penalties as predictions near true values to avoid overfitting.

### Model training strategy

Weighted edges between steroids were initialized based on their Spearman correlation coefficients, with edges from steroids to PI or SI set according to their correlations with CA. Edges from PI and SI to BA were intuitively set to 1 or -1, based on prior findings associating stress with aging acceleration and active sex hormone levels with youthfulness. Join nodes for intermediate steroids were processed through the activation function *ReLU*(*x*) = max (0, *x*), while other join nodes remained linear. The weights of all edges and biases were iteratively updated through backpropagation, managed by the Adam optimizer (*63*) with default momentum parameters *β*_*1*_ = 0.9, *β*_*2*_ = 0.999, and *ϵ* = 10^−8^.

### Hyperparameter tuning via cross-validation

Hyperparameters in DNN training, specifically the learning rate (*lr*) and epochs (*t*), were optimized using 5-fold cross-validation. The modeling group data was divided into five subsets, with each subset serving iteratively as the validation-fold while the remaining four formed the training-fold data. Optimal values for *lr* and *t* were determined by balancing four criteria: median validation loss, median validation loss, the difference in median loss between validation and training folds, standard deviation differences in loss between folds, and the training efficiency ratio (*t/lr*). This approach ensured robust model performance across a range of learning rates (*lr*: 0.001, 0.003, 0.005, 0.01) and epochs (*t*: 1000, 2000, 3000, 4000, 5000, 6000, 7000, 8000, 9000, 10000).

### Performance validation across sex and lifestyle factors

After training, the DNN model was transferred to an independent validation dataset to assess its performance. Loss values served as metrics to determine whether there were statistically significant performance differences between the training and validation datasets across sexes, using the Wilcoxon rank-sum test with Bonferroni correction and Kruskal-Wallis H test. The model’s performance was further evaluated using the defined angular difference *φ*, calculated as:

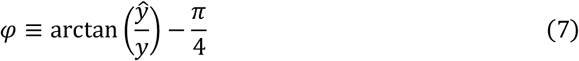

where *y* represents the true CA, and *ŷ* the predicted BA. We tested whether the median *φ* values for each group (modeling, smoking, and non-smoking) significantly differed from zero (*H*_*0*_: median = 0) using the Wilcoxon test with Bonferroni correction. Additionally, to examine the effects of smoking status, we assessed sex-specific differences in *φ* values using the Wilcoxon test, with Bonferroni correction applied. This analysis aimed to reveal how smoking habits may affect the model’s predictive capacity, providing insights into the potential impact of lifestyle factors on BA predictions.

### Sensitivity analysis of the DNN model

To evaluate the sensitivity of the DNN model to variations in steroid levels, each steroid concentration was independently increased by 100% within the dataset. For each steroid, the resulting change in predicted BA was calculated as a percentage difference from the baseline prediction. These calculations were performed across the entire modeling group, stratified by sex. Using these individual BA changes, we derived the mean change and 95% confidence intervals (CI), expressed as CI = *μ*_*Change*_ ± 1.96 × *σ*_*Change*_, to quantify each steroid’s impact on model predictions. An ANOVA test on the BA changes assessed significant sensitivity differences across steroids, followed by an effect size (*η*^2^) calculation. Post-hoc Tukey’s HSD analysis further identified steroids with distinct sensitivity effects.

## Acknowledgments

We thank Martin (WPI-ICReDD, Hokkaido Univ.) for DNN modeling advice, M. Okada for data correction guidance, Y. Pan for serum pre-experiment assistance, and S. Yashiro (VERITAS Corp.) for procuring serum samples.

## Author contributions

Conceptualization: QW, ZW

Funding acquisition: TT

Methodology: QW, ZW

Investigation: QW, ZW

Software: ZW

Formal analysis: ZW

Validation: QW, ZW

Visualization: QW, ZW

Supervision: KM, TT

Writing—original draft: QW, ZW

Writing—review & editing: QW, ZW, KM, TT

## Data and materials availability

All data needed to evaluate the conclusions in the paper are present in the paper and/or the Supplementary Materials. Raw data (under the CC-BY-NC License) and code (under the MIT License) for computational reproducibility are available in a public repository at Code Ocean (DOI: 10.24433/CO.3623382.v1).

## Supplementary Materials

### Supplementary Text

#### Preparation of standard stock, calibration, and quality control stock solutions

Each steroid standard was prepared in methanol at a concentration of 4 mg mL^-1^ as respective stock solutions and stored in -80 °C. These stock solutions were mixed and diluted to 10 ng μL^-1^ and with 40% MeOH as mixed stock solutions at -80 °C. The working standard solutions were prepared at concentrations of 0.4, 1, 2, 4, 10, 20, 40, 100, 160, 200, 400, 600, 1000, 1600, 2000, 8000 pg μL^-1^ with 40% MeOH for 16-OH-E1,7-OH-DHEA, TH-COL, 7-OH-P5, TH-COR, 11-OH-An, THB, APD, 17-OH-P5, 3β,5αTH-DOC, THS, 3α,5β-TH-DOC, COR, AT, COL, COB, COS, AE, E1, E2, DOC, TE, 17-OH-P4, DHT, and P4, and at concentrations of 0.8, 2, 4, 8, 20, 40, 80, 200, 320, 400, 800, 1200, 2000, 3200, 4000, 12000 pg μL^-1^ with 40% MeOH for DHEA, P5, E3, An and al-P5. An internal standard (IS) mixture solution of 0.1 ng μL^-1^ of 17-OH-P4-^13^C_3_, TE-^13^C_3_, AE-^13^C_3_, COR-^13^C_3_, P4-^13^C_3_, TH-COR-d_6_ and 16-OH-E1-^13^C_3_, and 0.2 ng μL^-1^ of COL-d_4_, COB-d_4_, E1-^13^C_3_, E3-d_3_ and 17-OH-P5-d_3_, and 0.4 ng μL^-1^ of DHEA-d_5_ and P5-^13^C_2_d_2_ was prepared in 40% MeOH.

For calibration and quality control samples, 24 μL of blank matrix (see below) was spiked with 6 μL of working standard solution and 4.8 μL of IS solution. The samples were evaporated to dryness and re-dissolved in 24 μL of 40% MeOH. Finally, the calibration samples were at levels of 0.01, 0.02, 0.05, 0.1, 0.2, 0.5, 1, 2, 5, 8, 10, 20, 30, 50, 80, 100, 300 pg μL^-1^ for 16-OH-E1,7-OH-DHEA, TH-COL, 7-OH-P5, TH-COR, 11-OH-An, THB, APD, 17-OH-P5, 3β,5αTH-DOC, THS, 3α,5β-TH-DOC, COR, AT, COL, COB, COS, AE, E1, E2, DOC, TE, 17-OH-P4, DHT, and P4, and at levels of 0.02, 0.04, 0.1, 0.2, 0.4, 1, 2, 4, 10, 16, 20, 30, 50, 100, 160, 200, 600 pg μL^-1^ for DHEA, P5, E3, An and al-P5.

#### Preparation of blood sample blank matrix for LC-MS/MS validation

For method validation, charcoal was added during the sample preparation procedure. Serum samples were protein precipitated with ACN. After centrifuging, the supernatant was collected and diluted with H_2_O until 10% ACN. Then charcoal was added to strip the steroids from the system. 0.6 mg of activated charcoal was added per microliter of serum, followed by vortexing and centrifugation. The supernatant was loaded on the bond elute column for further purification. The eluate was evaporated to dryness with speed-vac and re-dissolved with 40% MeOH to form 10 μL·μL^-1^ blank matrix.

Since the concentration of sex steroid hormones are extremely low in the serum of very old people, researchers used such serum as sex steroid free blank matrix. However, the corticosteroids play important roles human body regardless of age, it was impossible to find the serum samples for which the corticosteroid did not exist. In this case, charcoal extraction was used for the blank matrix preparation procedure. In should be noted that charcoal might strip out the interfering compounds, which might be the weakness of using a blank matrix prepared in this way.

#### LC-MS/MS method validation

The developed method was satisfactorily validated in terms of the LOQ, linear range, extraction recovery, precision, and accuracy.

### Calibration curves and LOQ

The calibration curves, correlation coefficients, linear ranges, and LOQs of the 13 steroids in the spiked blank matrix are shown in Table S4. The calibration curve was constructed using the peak area ratios of a compound to IS versus the ratios of concentrations of a compound at different levels to the concentration of IS. The correlation coefficient square (r^2^) was calculated. LOQ was tested at a signal to noise (S/N) of 10. Good linearity was observed for all 29 steroids within the ranges (0.01-10 pg µL^-1^ for 7-OH-DHEA, 0.02-2 pg µL^-1^ for 3β,5α-TH-DOC, 0.02-5 pg µL^-1^ for 16-OH-E1, 0.02-20 pg µL^-1^ for TH-COR, 0.04-10 pg µL^-1^ for DOC, 0.05-10 pg µL^-1^ for AT, 0.05-20 pg µL^-1^ for COS, 0.05-50 pg µL^-1^ for 7-OH-P5, and TH-COL, 0.1-8 pg µL^-1^ for THB, 0.1-20 pg µL^-1^ for 17-OH-P4 and P4, 0.1-50 pg µL^-1^ for 11-OH-An and 17-OH-P5, 0.2-20 pg µL^-1^ for APD, THS, 3α,5β-TH-DOC, COB, AE, E1, E2, TE, DHT and An, 0.4-100 pg µL^-1^ for P5, 0.5-50 pg µL^-1^ for COR, 1-100 pg µL^-1^ for DHEA, E3 and al-P5, and 2-300 pg µL^-1^ for COL. The linear correlation coefficient square (r^2^) was greater than 0.9921. The lower LOQ (LLOQ) was 0.005-0.288 pg µL^-1^ for serum, suggesting that the developed method is highly sensitive for the quantification of the steroids.

#### Matrix effect and recovery in LC-MS/MS analysis

To evaluate the matrix effect (ME) and recovery (R), low (0.05, 0.2, 0.4, 1, 2, 5, 10 or 50 pg µL^-1^), medium (0.2, 0.4, 1, 2, 5, 10, 20 or 80 pg µL^-1^) and high (1, 2, 5, 10, 20, 40 or 100 pg µL^-1^) concentrations of spiked blank matrix and serum samples (n = 3) were assessed. The matrix effect value was calculated as ME (%) = A_matrix_/A_solution_ × 100, where A_solution_ is the compound peak area of 10 μL of pure standard and A_matrix_ is the compound peak area of blank matrix spiked with 10 μL standard sample. The recovery value was calculated as R (%) = A_pre-spike_/A_post-spike_ × 100, where A_pre-spike_ is the compound peak area of serum spiked with 10 μL standards before extraction and A_post-spike_ is the compound peak area of serum spiked with 10 μL standards after extraction. The extraction matrix effect and recoveries (%) were within the range of 88.09-127.64.40% and 75.15-121.00% in Table S5.

#### Accuracy and precision in LC-MS/MS analysis

The precision and accuracy of the method were assessed by performing six replicates of serum samples spiked with low (0.05, 0.2, 0.4, 1, 2, 5, 10 or 50 pg µL^-1^), medium (0.2, 0.4, 1, 2, 5, 10, 20 or 80 pg µL^-1^) and high (1, 2, 5, 10, 20, 40 or 100 pg µL^-1^) of steroids (Table S6). Accuracy was calculated as the averaged percentage for the measured concentrations to the real concentrations. Precision was expressed as the RSDs (relative standard deviation) of the measured concentrations and was performed on three separate days. The low and high concentration of steroids spiked in the intra- and inter-batch samples were obtained within acceptable ranges. Table S6 summarizes the accuracy and precision data for serum. The results indicated that the method shows a moderately good precision and accuracy. The accuracy (%) was 74.75-113.05 while the intra-day precision (%) was 0.002-17.65 and the Inter-day precision (%) was 0.001-23.85. The method was considered to be suitable in terms of accuracy and precision.

**Fig. S1.**
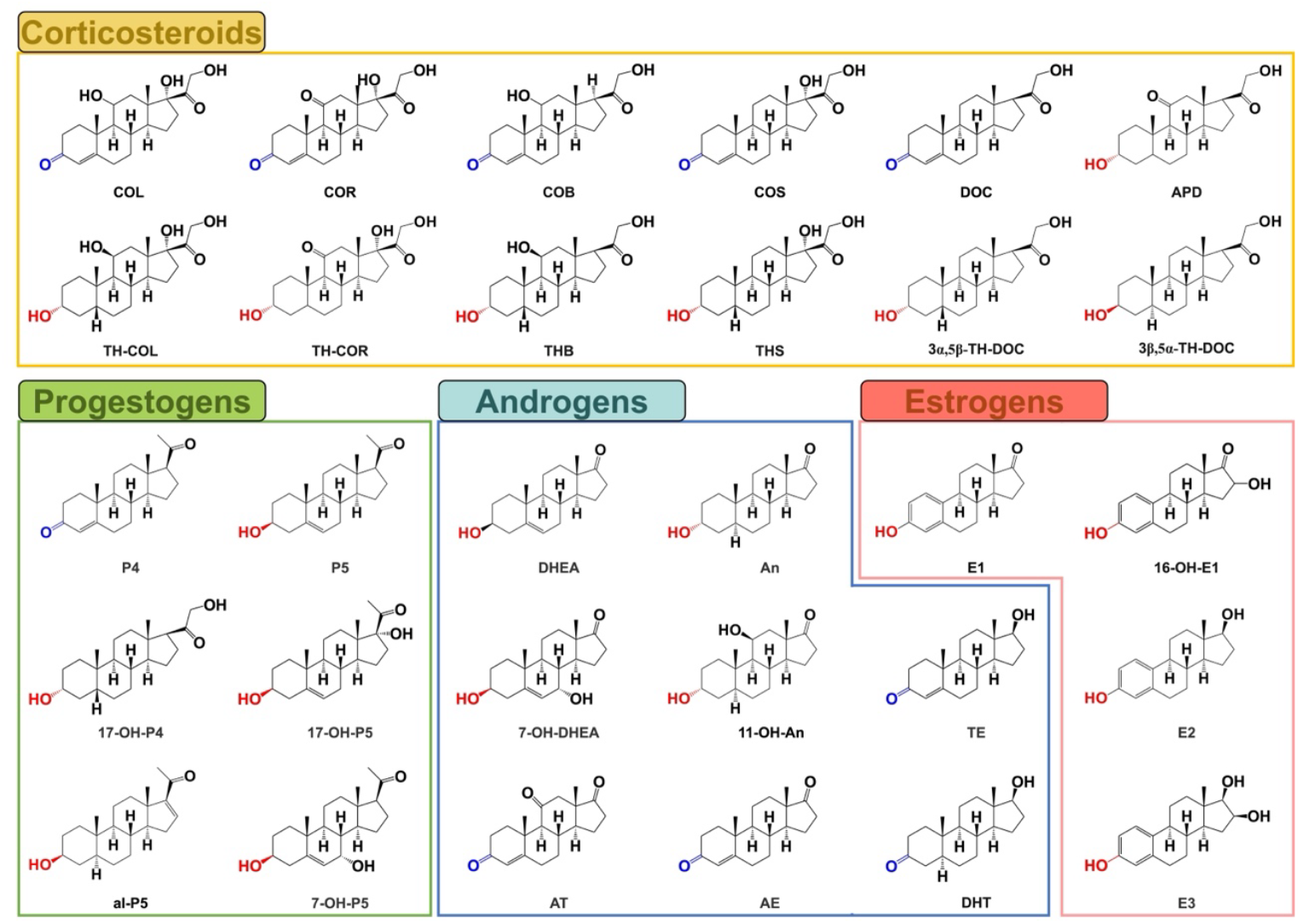
Structures of steroids analyzed in this study. The steroid structures are categorized by color-coded boxes: yellow for corticosteroids, pink for estrogens, green for progestogens, and blue for androgens. 11-OH-An, 11-β-hydroxyandrosterone; 16-OH-E1, 16-hydroxyestrone; 17-OH-P5, 17α-hydroxypregnenolone; 17-OH-P4, 17α-Hydroxyprogesterone; 7-OH-DHEA, 7α-hydroxydehydroepiandrosterone; 7-OH-P5, 7α-hydroxypregnenolone; Androstenedione, AE; al-P5, allopregnenolone; An, androsterone; APD, alphadolone; AT, Adrenosterone; COB, Corticosterone; COL, Cortisol; COR, cortisone; COS, 11-Deoxycortisol; DHEA, dehydroepiandrosterone; DHT, dihydrotestosterone; DOC, 11-Deoxycorticosterone; E1, Estrone; E2, Estradiol; E3, estriol; P4, Progesterone; P5, pregnenolone; TE, Testosterone; THB, tetrahydrocorticosterone; TH-COL, tetrahydrocortisol; TH-COR, tetrahydrocortisone; TH-DOC, 3β,5α-tetrahydrodeoxycorticosterone; THS, tetrahydrodeoxycortisol.

**Fig. S2.**
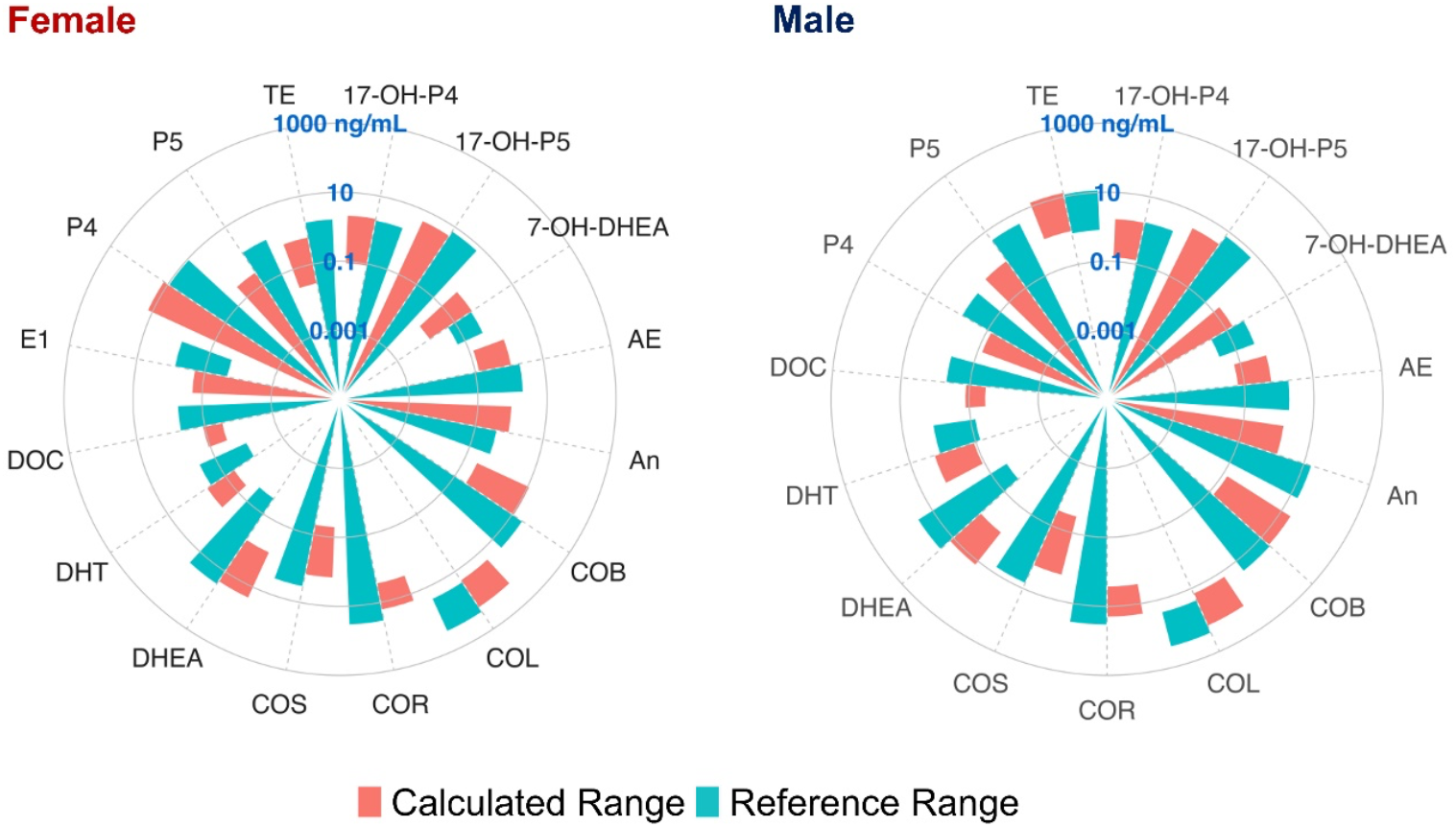
Comparison of steroid concentrations with published reference ranges. Radar plots compare steroid concentrations measured by the developed method (red bars) with reference ranges from the literature (blue bars) for both male and female healthy subjects. Detailed reference ranges are summarized in Table S5.

**Fig. S3.**
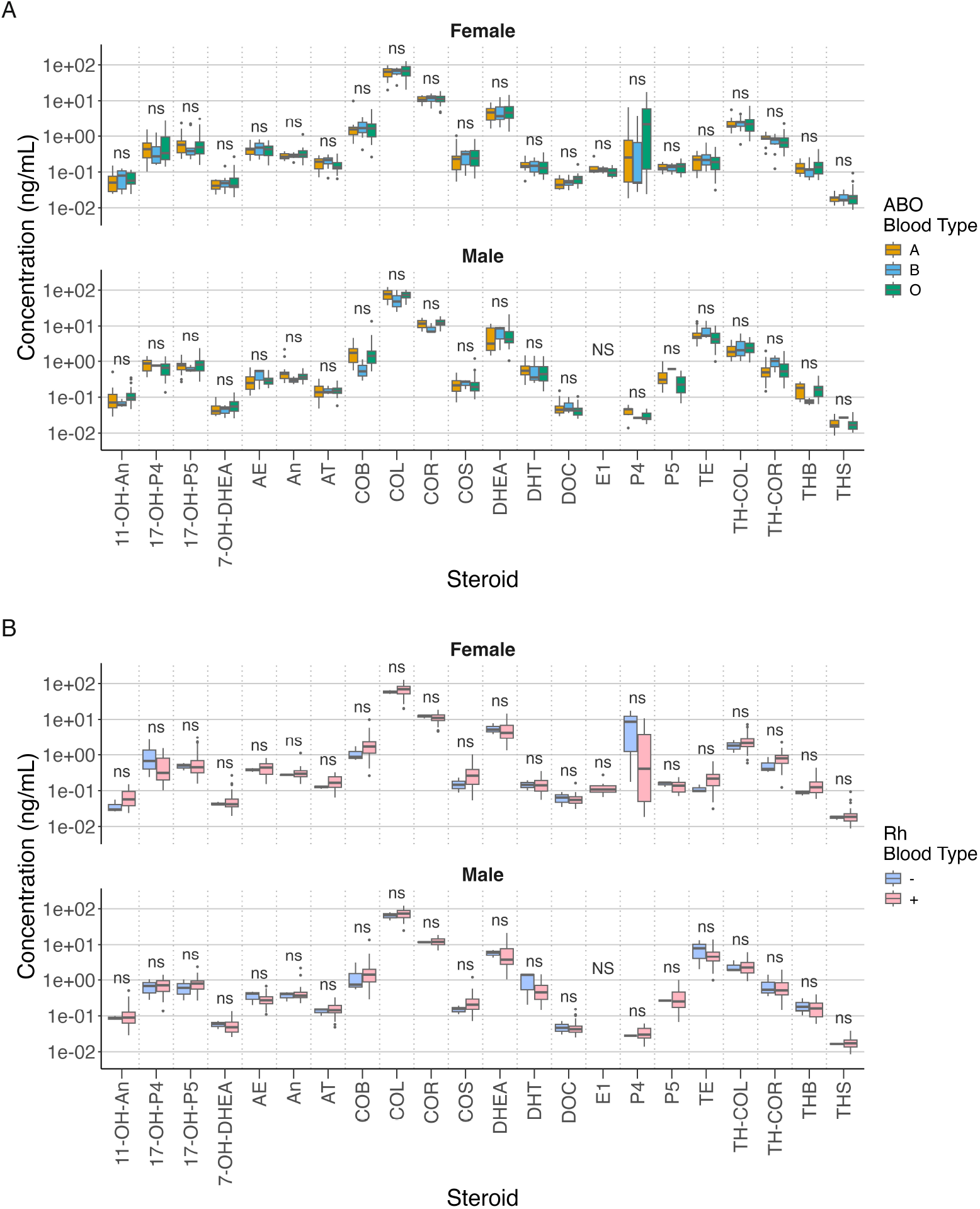
Distribution of steroid concentrations before and after scaling for different blood types. Univariate analysis for (**A**) ABO blood type and (**B**) Rh blood type was performed using the Kruskal-Wallis test with Bonferroni correction. The data for ABO blood type include females (A: n = 11; B: n = 11; O: n = 26; AB: n = 1, excluded from analysis) and males (A: n = 16; B: n = 3; O: n = 30) from the modeling dataset. The Rh blood type data include females (+: n = 46; -: n = 3) and males (+: n = 46; -: n = 3) also derived from the modeling dataset. Statistical significance, NS, all zero values (non-test); ns, P_adj_ ≥ 0.05.

**Fig. S4.**
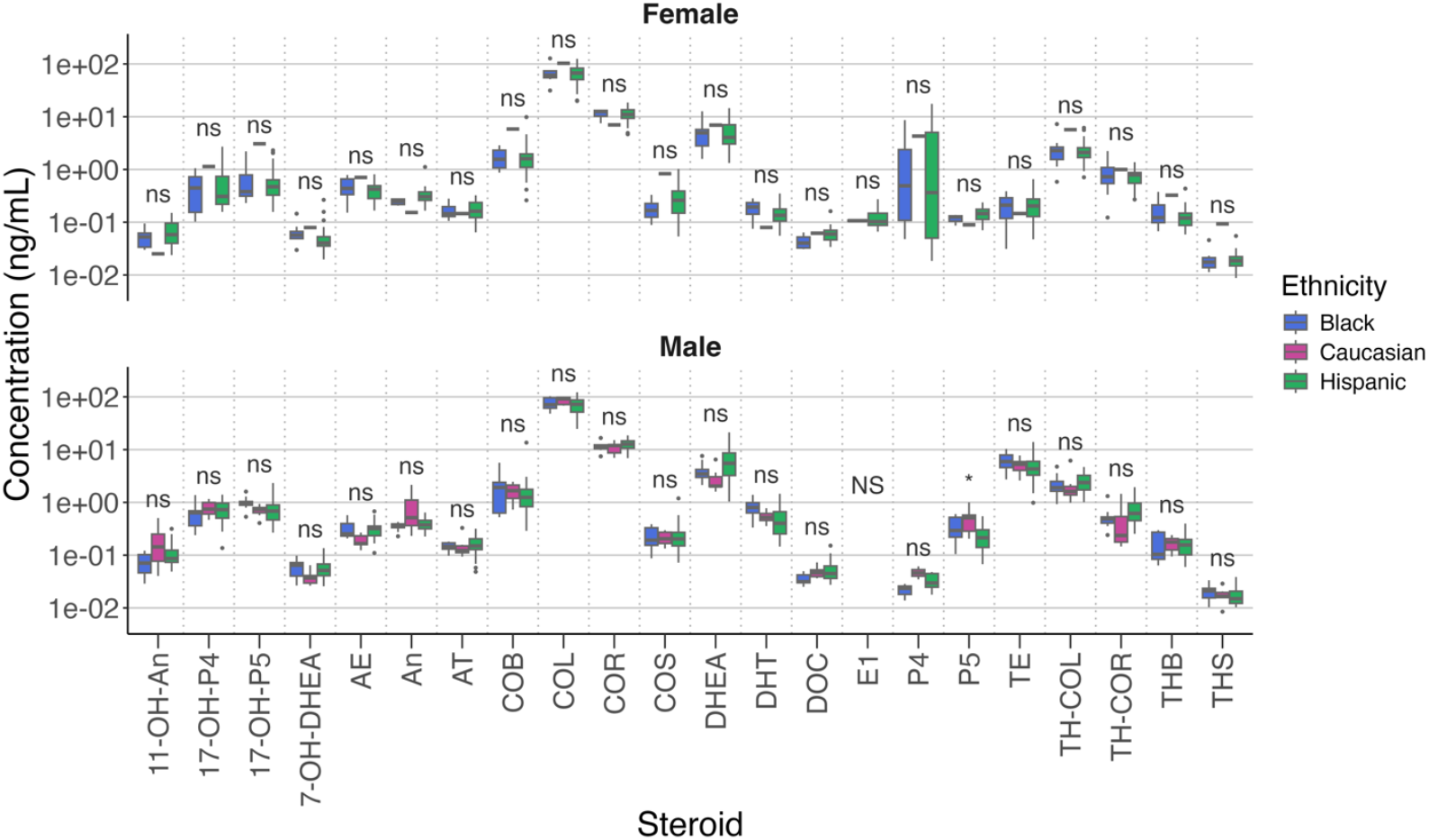
Distribution of steroid concentrations before and after scaling for different ethnicities. Univariate analysis for ethnicity, using the Kruskal-Wallis test adjusted by Bonferroni correction. The data for ethnicity include females (Black: n = 8; Caucasian: n = 1; Hispanic: n = 40) and males (Black: n = 9; Caucasian: n = 7; Hispanic: n = 33) from the modeling dataset. Statistical significance, NS, all zero values (non-test); ns, P_adj_ ≥ 0.05; * P_adj_ < 0.05.

**Fig. S5.**
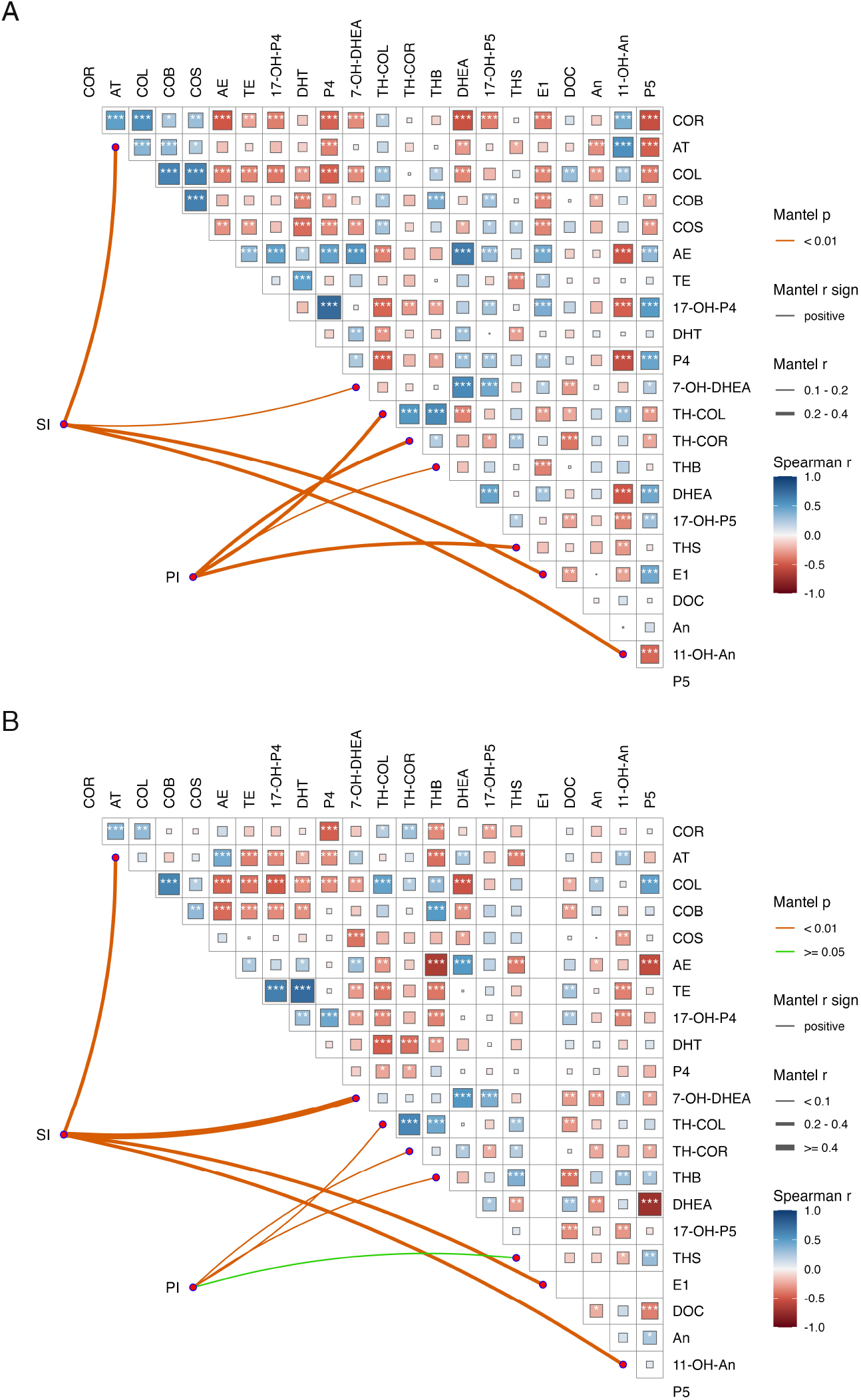
Initial DNN edge weights based on steroid and CA correlations. Spearman correlation heatmaps displaying inter-steroid correlations, along with associated p-values, for (**A**) females and (**B**) males. Correlations between CA and eight steroids relevant to PI and SI are represented by connecting lines, with line weights indicating correlation strength and colors indicating significance levels (p-values). Statistical significance, blank, all zero values (non-test); * P_adj_ < 0.05; ** P_adj_ < 0.01; *** P_adj_ < 0.001.

**Fig. S6.**
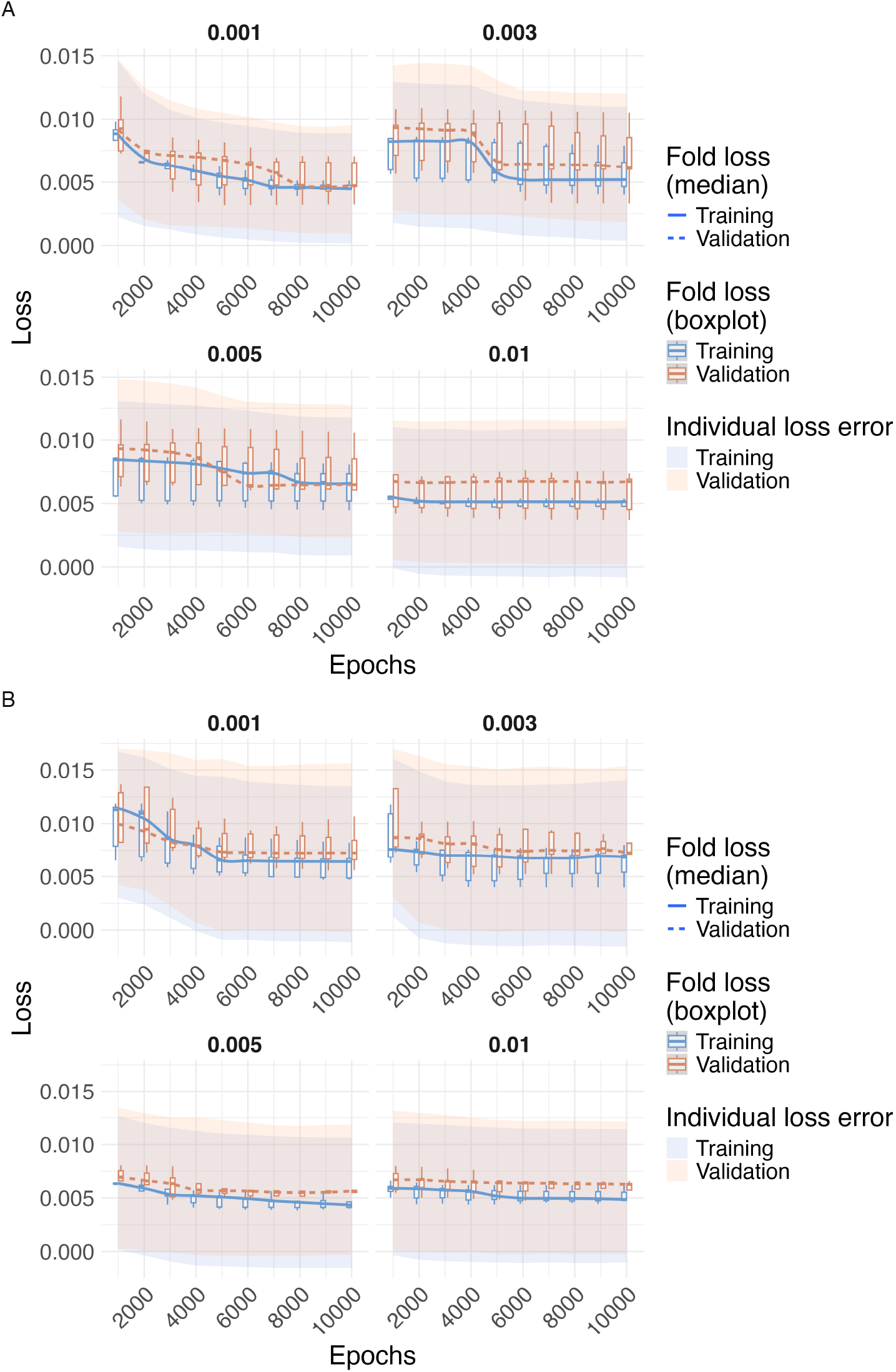
Hyperparameter optimization through 5-fold cross-validation. Model loss distributions across epochs for different learning rates in both (**A**) female and (**B**) male models. Each subplot represents model losses on the training (modeling) and validation folds at intervals of 1000 epochs, up to a maximum of 10000 epochs. The learning rate value for each subplot is indicated in the respective panel.

**Fig. S7.**
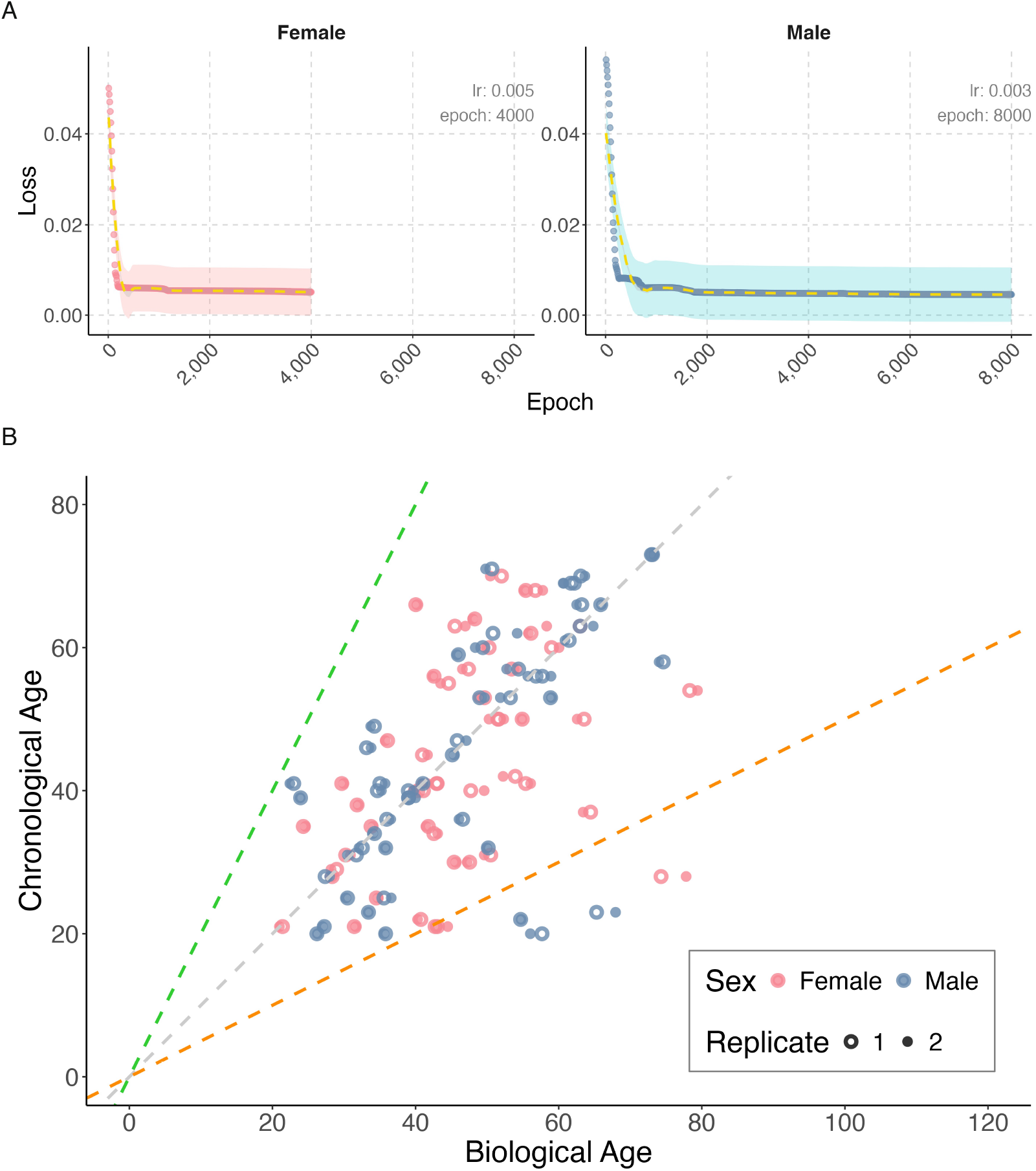
Training performance of the DNN model with optimal hyperparameters. Training curves for (**A**) females and (**B**) males, illustrating performance metrics based on the corresponding optimal learning rates and epochs. Shaded areas represent the standard deviation of individual training losses. (**C**) Scatter plot comparing predicted BA against true CA for females (n = 98, including two replicates) and males (n = 98, including two replicates). lr, learning rate.

**Fig. S8.**
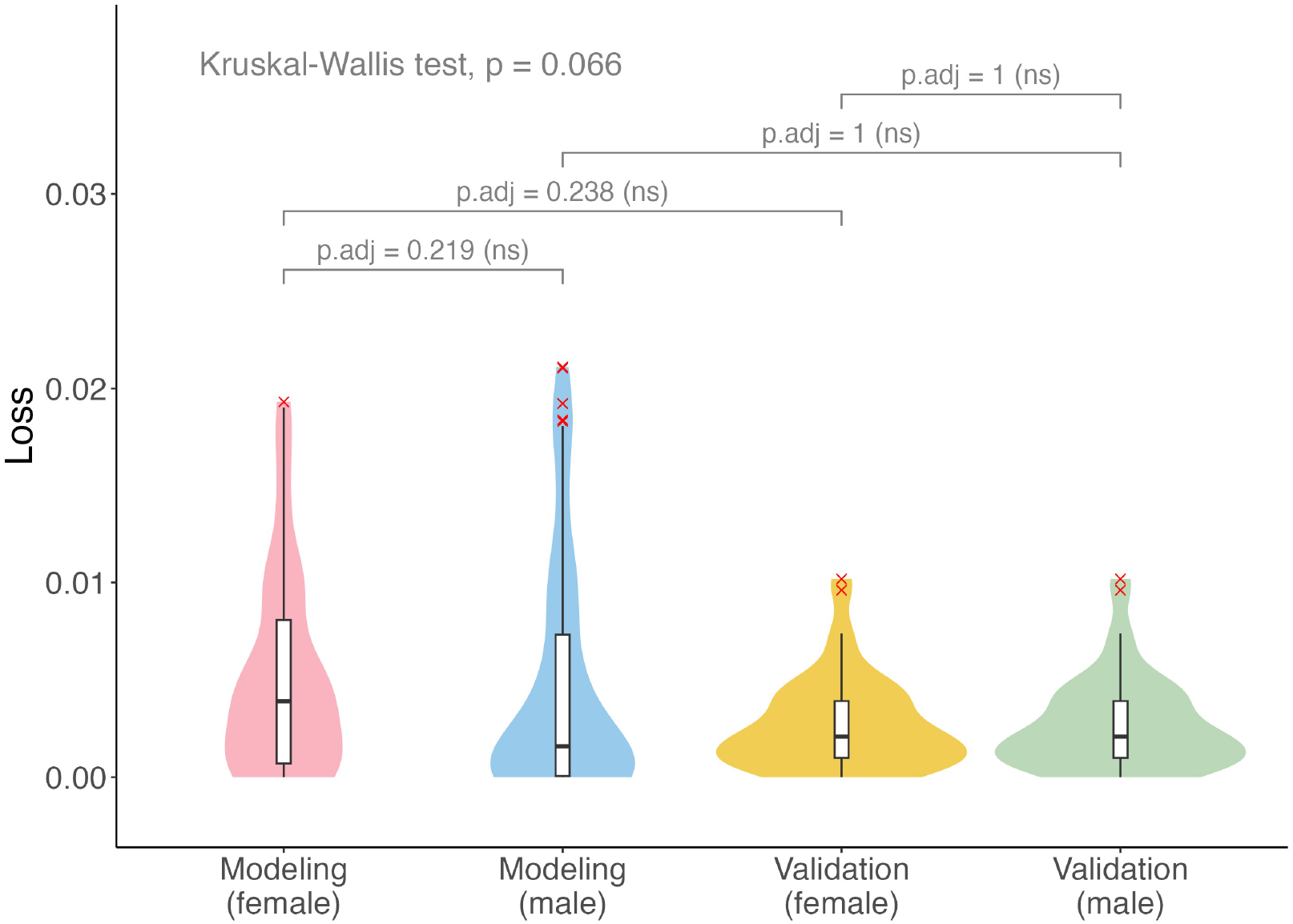
Statistical analysis of WSATL values across all groups. WSATL values were statistically analyzed across all groups using the Kruskal-Wallis test with Bonferroni correction for overall comparisons, and the Wilcoxon test with Bonferroni adjustment for pairwise comparisons between specific groups. The modeling dataset includes females (n = 98, with two replicates) and males (n = 98, with two replicates), while the validation dataset comprises females (n = 50, with two replicates) and males (n = 50, with two replicates). Statistical significance, ns, P_adj_ ≥ 0.05.

**Fig. S9.**
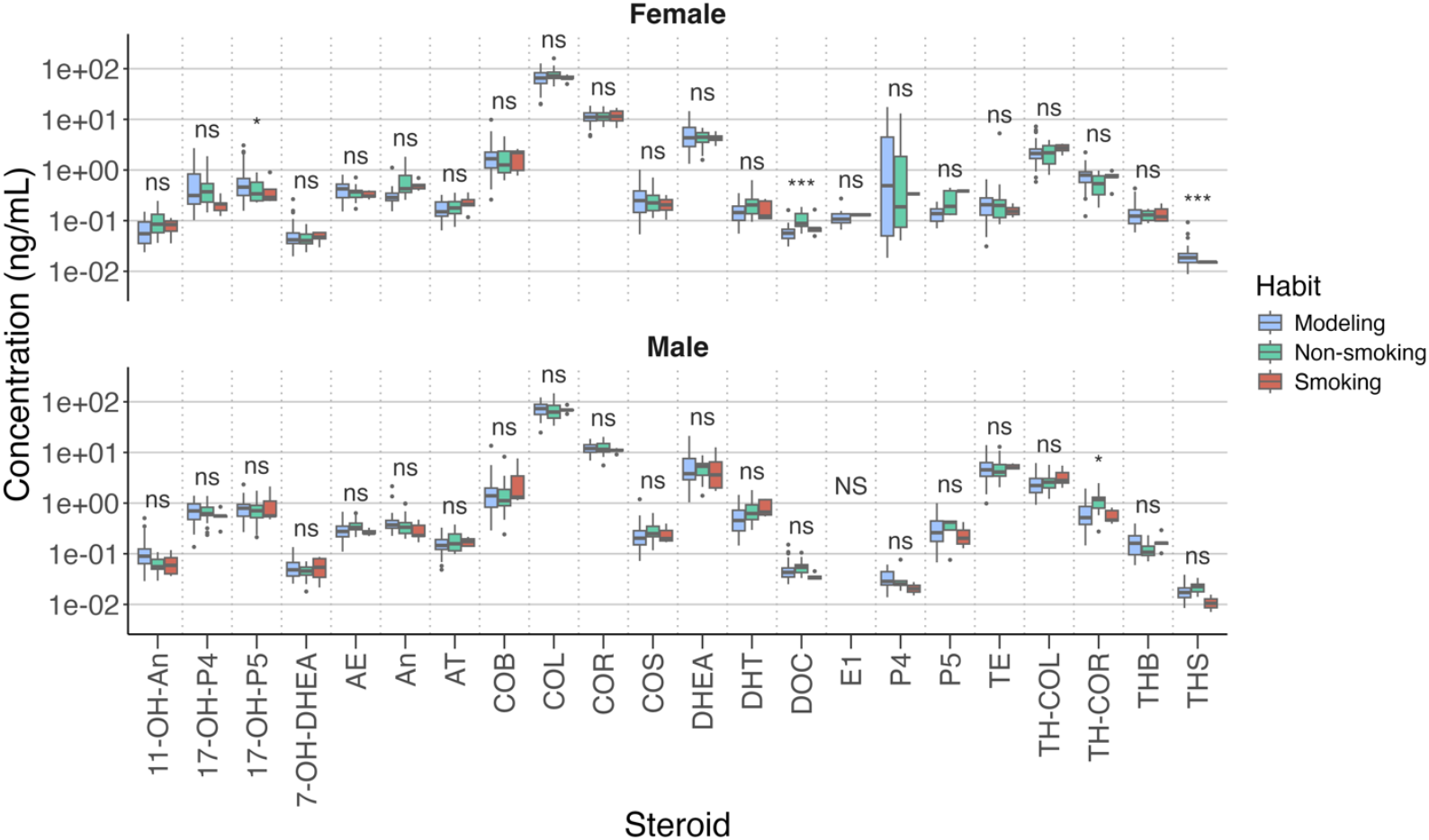
Distribution of steroid concentrations before and after scaling for different smoking habit. Univariate analysis for smoking habit, using the Kruskal-Wallis test adjusted by Bonferroni correction. The data for smoking habit include females (Modeling, habit unknown: n = 49; Non-smoking: n = 20; Smoking: n = 5) and males (Modeling, habit unknown: n = 49; Non-smoking: n = 20; Smoking: n = 5) from both the modeling and validation dataset. Statistical significance, NS, all zero values (non-test); ns, P_adj_ ≥ 0.05; * P_adj_ < 0.05; ** P_adj_ < 0.01; *** P_adj_ < 0.001.

## References

1. L. Fontana, L. Partridge, V. D. Longo, Extending healthy life span—from yeast to humans. science 328, 321–326 (2010).

2. A. Picca, H. J. Coelho-Junior, R. Calvani, E. Marzetti, D. L. Vetrano, Biomarkers shared by frailty and sarcopenia in older adults: A systematic review and meta-analysis. Ageing research reviews 73, 101530 (2022).

3. V. R. Varma, A. M. Oommen, S. Varma, R. Casanova, Y. An, R. M. Andrews, R. O’Brien, O. Pletnikova, J. C. Troncoso, J. Toledo, R. Baillie, M. Arnold, G. Kastenmueller, K. Nho, P. M. Doraiswamy, A. J. Saykin, R. Kaddurah-Daouk, C. Legido-Quigley, M. Thambisetty, Brain and blood metabolite signatures of pathology and progression in Alzheimer disease: A targeted metabolomics study. PLoS Med 15, e1002482 (2018).

4. L. Feng, J. Li, R. Zhang, Current research status of blood biomarkers in Alzheimer’s disease: Diagnosis and prognosis. Ageing Research Reviews 72, 101492 (2021).

5. M. Rodriguez, C. Rodriguez-Sabate, I. Morales, A. Sanchez, M. Sabate, Parkinson’s disease as a result of aging. Aging cell 14, 293–308 (2015).

6. Z. Wang, L. Bian, C. Mo, H. Shen, L. J. Zhao, K. J. Su, M. Kukula, J. T. Lee, D. W. Armstrong, R. Recker, J. Lappe, L. F. Bonewald, H. W. Deng, M. Brotto, Quantification of aminobutyric acids and their clinical applications as biomarkers for osteoporosis. Commun Biol 3, 39 (2020).

7. C. E. Conti Filho, C. Marcolongo-Pereira, J. V. Rossoni Junior, R. M. Barcelos, O. Chiarelli-Neto, F. C. d. A. Q. Castro, S. F. Teixeira, N. J. Mezzomo, Advances in Alzheimer’s disease’s pharmacological treatment. Frontiers in Pharmacology 14, 1101452 (2023).

8. P. Sivanandy, T. C. Leey, T. C. Xiang, T. C. Ling, S. A. Wey Han, S. L. A. Semilan, P. K. Hong, Systematic review on Parkinson’s disease medications, emphasizing on three recently approved drugs to control Parkinson’s symptoms. International journal of environmental research and public health 19, 364 (2021).

9. Y. Rolland, C. Dray, B. Vellas, P. D. S. Barreto, Current and investigational medications for the treatment of sarcopenia. Metabolism, 155597 (2023).

10. I. Foessl, H. P. Dimai, B. Obermayer-Pietsch, Long-term and sequential treatment for osteoporosis. Nature Reviews Endocrinology 19, 520–533 (2023).

11. J. Zierer, C. Menni, G. Kastenmüller, T. D. Spector, Integration of ‘omics’ data in aging research: from biomarkers to systems biology. Aging cell 14, 933–944 (2015).

12. F. Galkin, P. Mamoshina, A. Aliper, J. P. de Magalhaes, V. N. Gladyshev, A. Zhavoronkov, Biohorology and biomarkers of aging: Current state-of-the-art, challenges and opportunities. Ageing Res Rev 60, 101050 (2020).

13. B. Osborne, D. Bakula, M. Ben Ezra, C. Dresen, E. Hartmann, S. M. Kristensen, G. V. Mkrtchyan, M. H. Nielsen, M. A. Petr, M. Scheibye-Knudsen, New methodologies in ageing research. Ageing Res Rev 62, 101094 (2020).

14. A. M. Herskind, M. McGue, N. V. Holm, T. I. Sörensen, B. Harvald, J. W. Vaupel, The heritability of human longevity: a population-based study of 2872 Danish twin pairs born 1870–1900. Human genetics 97, 319–323 (1996).

15. S. Song, Y. Stern, Y. Gu, Modifiable lifestyle factors and cognitive reserve: a systemic review of current evidence. Ageing research reviews, 101551 (2021).

16. E. Nakamura, A study on the basic nature of human biological aging processes based upon a hierarchical factor solution of the age-related physiological variables. Mechanisms of ageing and development 60, 153–170 (1991).

17. M. E. Levine, Modeling the rate of senescence: can estimated biological age predict mortality more accurately than chronological age? Journals of Gerontology Series A: Biomedical Sciences and Medical Sciences 68, 667–674 (2013).

18. A. Comfort, Test-battery to measure ageing-rate in man. The Lancet 294, 1411–1415 (1969).

19. T. Kirkwood, Alex Comfort and the measure of aging. Experimental gerontology 33, 135–140 (1998).

20. S. H. Jackson, M. R. Weale, R. A. Weale, Biological age—what is it and can it be measured? Archives of gerontology and geriatrics 36, 103–115 (2003).

21. J. B. Dowd, N. Goldman, Do biomarkers of stress mediate the relation between socioeconomic status and health? Journal of Epidemiology & Community Health 60, 633–639 (2006).

22. T. E. Johnson, Recent results: biomarkers of aging. Experimental gerontology 41, 1243–1246 (2006).

23. L. Hayflick, Biological aging is no longer an unsolved problem. Annals of the New York academy of Sciences 1100, 1–13 (2007).

24. J. Bortz, A. Guariglia, L. Klaric, D. Tang, P. Ward, M. Geer, M. Chadeau-Hyam, D. Vuckovic, P. K. Joshi, Biological age estimation using circulating blood biomarkers. Communications biology 6, 1089 (2023).

25. L. Wu, X. Xie, T. Liang, J. Ma, L. Yang, J. Yang, L. Li, Y. Xi, H. Li, J. Zhang, Integrated multi-omics for novel aging biomarkers and antiaging targets. Biomolecules 12, 39 (2021).

26. S. Pal, J. K. Tyler, Epigenetics and aging. Science advances 2, e1600584 (2016).

27. S. Fiacco, A. Walther, U. Ehlert, Steroid secretion in healthy aging. Psychoneuroendocrinology 105, 64–78 (2019).

28. M. C. Velarde, Mitochondrial and sex steroid hormone crosstalk during aging. Longevity & healthspan 3, 1–10 (2014).

29. S. Kashif Zaidi, W.-J. Shen, S. Azhar, Impact of aging on steroid hormone biosynthesis and secretion. Open Longevity Science 6, (2012).

30. D. J. Panyard, B. Yu, M. P. Snyder, The metabolomics of human aging: Advances, challenges, and opportunities. Science Advances 8, eadd6155 (2022).

31. J. Rutledge, H. Oh, T. Wyss-Coray, Measuring biological age using omics data. Nature Reviews Genetics 23, 715–727 (2022).

32. B. Lehallier, D. Gate, N. Schaum, T. Nanasi, S. E. Lee, H. Yousef, P. Moran Losada, D. Berdnik, A. Keller, J. Verghese, S. Sathyan, C. Franceschi, S. Milman, N. Barzilai, T. Wyss-Coray, Undulating changes in human plasma proteome profiles across the lifespan. Nat Med 25, 1843–1850 (2019).

33. M. J. Peters, R. Joehanes, L. C. Pilling, C. Schurmann, K. N. Conneely, J. Powell, E. Reinmaa, G. L. Sutphin, A. Zhernakova, K. Schramm, The transcriptional landscape of age in human peripheral blood. Nature communications 6, 1–14 (2015).

34. S. Horvath, DNA methylation age of human tissues and cell types. Genome biology 14, 1–20 (2013).

35. J. Kristic, F. Vuckovic, C. Menni, L. Klaric, T. Keser, I. Beceheli, M. Pucic-Bakovic, M. Novokmet, M. Mangino, K. Thaqi, P. Rudan, N. Novokmet, J. Sarac, S. Missoni, I. Kolcic, O. Polasek, I. Rudan, H. Campbell, C. Hayward, Y. Aulchenko, A. Valdes, J. F. Wilson, O. Gornik, D. Primorac, V. Zoldos, T. Spector, G. Lauc, Glycans are a novel biomarker of chronological and biological ages. J Gerontol A Biol Sci Med Sci 69, 779–789 (2014).

36. J. Hertel, N. Friedrich, K. Wittfeld, M. Pietzner, K. Budde, S. Van der Auwera, T. Lohmann, A. Teumer, H. Volzke, M. Nauck, H. J. Grabe, Measuring Biological Age via Metabonomics: The Metabolic Age Score. J Proteome Res 15, 400–410 (2016).

37. X. Ni, H. Zhao, R. Li, H. Su, J. Jiao, Z. Yang, Y. Lv, G. Pang, M. Sun, C. Hu, Development of a model for the prediction of biological age. Computer Methods and Programs in Biomedicine 240, 107686 (2023).

38. B. Couvy-Duchesne, J. Faouzi, B. Martin, E. Thibeau–Sutre, A. Wild, M. Ansart, S. Durrleman, D. Dormont, N. Burgos, O. Colliot, Ensemble learning of convolutional neural network, support vector machine, and best linear unbiased predictor for brain age prediction: Aramis contribution to the predictive analytics competition 2019 challenge. Frontiers in Psychiatry 11, 593336 (2020).

39. L. Sagers, L. Melas-Kyriazi, C. J. Patel, A. K. Manrai, Prediction of chronological and biological age from laboratory data. Aging (Albany NY) 12, 7626 (2020).

40. M. E. Levine, Modeling the rate of senescence: can estimated biological age predict mortality more accurately than chronological age? J Gerontol A Biol Sci Med Sci 68, 667–674 (2013).

41. P. Mamoshina, K. Kochetov, F. Cortese, A. Kovalchuk, A. Aliper, E. Putin, M. Scheibye-Knudsen, C. R. Cantor, N. M. Skjodt, O. Kovalchuk, A. Zhavoronkov, Blood Biochemistry Analysis to Detect Smoking Status and Quantify Accelerated Aging in Smokers. Sci Rep 9, 142 (2019).

42. E. Putin, P. Mamoshina, A. Aliper, M. Korzinkin, A. Moskalev, A. Kolosov, A. Ostrovskiy, C. Cantor, J. Vijg, A. Zhavoronkov, Deep biomarkers of human aging: application of deep neural networks to biomarker development. Aging (Albany NY) 8, 1021 (2016).

43. P. Mamoshina, K. Kochetov, E. Putin, F. Cortese, A. Aliper, W. S. Lee, S. M. Ahn, L. Uhn, N. Skjodt, O. Kovalchuk, M. Scheibye-Knudsen, A. Zhavoronkov, Population Specific Biomarkers of Human Aging: A Big Data Study Using South Korean, Canadian, and Eastern European Patient Populations. J Gerontol A Biol Sci Med Sci 73, 1482–1490 (2018).

44. B. Lonnqvist, A. Bornet, A. Doerig, M. H. Herzog, A comparative biology approach to DNN modeling of vision: A focus on differences, not similarities. Journal of vision 21, 17–17 (2021).

45. Z.-Y. Zhang, X.-R. Sheng, Y. Zhang, B. Jiang, S. Han, H. Deng, B. Zheng, in Proceedings of the 31st ACM international conference on information & knowledge management. (2022), pp. 2671–2680.

46. N. Sapoval, A. Aghazadeh, M. G. Nute, D. A. Antunes, A. Balaji, R. Baraniuk, C. Barberan, R. Dannenfelser, C. Dun, M. Edrisi, Current progress and open challenges for applying deep learning across the biosciences. Nature Communications 13, 1728 (2022).

47. L. Ferrucci, G. A. Kuchel, Heterogeneity of aging: individual risk factors, mechanisms, patient priorities, and outcomes. Journal of the American Geriatrics Society 69, 610 (2021).

48. A. Thalén, A. Ledberg, Consequences of heterogeneity in aging: parental age at death predicts midlife all-cause mortality and hospitalization in a Swedish national birth cohort. BMC geriatrics 24, 207 (2024).

49. Q. Wang, K. Shimizu, K. Maehata, Y. Pan, K. Sakurai, T. Hikida, Y. Fukada, T. Takao, Lithium ion adduction enables UPLC-MS/MS-based analysis of multi-class 3-hydroxyl group-containing keto-steroids [S]. Journal of lipid research 61, 570–579 (2020).

50. S. Woloshin, L. M. Schwartz, H. G. Welch, The risk of death by age, sex, and smoking status in the United States: putting health risks in context. Journal of the National Cancer Institute 100, 845–853 (2008).

51. J. Coste, L. Quinquis, S. d’Almeida, E. Audureau, Smoking and health-related quality of life in the general population. Independent relationships and large differences according to patterns and quantity of smoking and to gender. PloS one 9, e91562 (2014).

52. M. Turton, T. Deegan, Circadian variations of plasma catecholamine, cortisol and immunoreactive insulin concentrations in supine subjects. Clinica Chimica Acta 55, 389–397 (1974).

53. E. Van Cauter, R. Leproult, D. J. Kupfer, Effects of gender and age on the levels and circadian rhythmicity of plasma cortisol. The Journal of Clinical Endocrinology & Metabolism 81, 2468–2473 (1996).

54. E. T. Klopack, J. E. Carroll, S. W. Cole, T. E. Seeman, E. M. Crimmins, Lifetime exposure to smoking, epigenetic aging, and morbidity and mortality in older adults. Clinical Epigenetics 14, 72 (2022).

55. J. S. Tolstrup, U. A. Hvidtfeldt, E. M. Flachs, D. Spiegelman, B. L. Heitmann, K. Bälter, U. Goldbourt, G. Hallmans, P. Knekt, S. Liu, Smoking and risk of coronary heart disease in younger, middle-aged, and older adults. American journal of public health 104, 96–102 (2014).

56. X. Zhu, J. Xue, R. Maimaitituerxun, H. Xu, Q. Zhou, Q. Zhou, W. Dai, W. Chen, Relationship between dietary macronutrients intake and biological aging: a cross-sectional analysis of NHANES data. European Journal of Nutrition 63, 243–251 (2024).

57. L. Oblak, J. van der Zaag, A. T. Higgins-Chen, M. E. Levine, M. P. Boks, A systematic review of biological, social and environmental factors associated with epigenetic clock acceleration. Ageing research reviews 69, 101348 (2021).

58. J. K. Kresovich, A. M. Martinez Lopez, E. L. Garval, Z. Xu, A. J. White, D. P. Sandler, J. A. Taylor, Alcohol consumption and methylation-based measures of biological age. The Journals of Gerontology: Series A 76, 2107–2111 (2021).

59. S. R. Beach, M. V. Dogan, M. K. Lei, C. E. Cutrona, M. Gerrard, F. X. Gibbons, R. L. Simons, G. H. Brody, R. A. Philibert, Methylomic aging as a window onto the influence of lifestyle: tobacco and alcohol use alter the rate of biological aging. Journal of the American geriatrics society 63, 2519–2525 (2015).

60. I. K. Yeo, R. A. Johnson, A new family of power transformations to improve normality or symmetry. Biometrika 87, 954–959 (2000).

61. M. Kanehisa, Toward understanding the origin and evolution of cellular organisms. Protein Science 28, 1947–1951 (2019).

62. M. Kanehisa, S. Goto, KEGG: kyoto encyclopedia of genes and genomes. Nucleic acids research 28, 27–30 (2000).

63. D. P. Kingma, Adam: A method for stochastic optimization. arXiv preprint arXiv:1412.6980, (2014).

